# A Preoptic Neuronal Population Regulates Energy Expenditure and Balance

**DOI:** 10.64898/2026.04.09.715318

**Authors:** Juan Liu, Aaron L. Cone, Daniel Ferguson, Amanda Breese, Hannah E Skelton, Abraham Escobedo, Alexxai V. Kravitz, Eric C. Landsness, Brian N. Finck, Aaron J. Norris

## Abstract

Maintaining energy balance requires coordination between food intake and energy expenditure, yet the neural pathways that regulate energy expenditure remain unclear. This study identifies kappa opioid receptor–expressing neurons in the preoptic area of the hypothalamus as a key regulator of whole-body metabolism. Using mouse models combined with fiber photometry, chemogenetic activation and inhibition, and chronic disruption of synaptic output, the results show that activity of these neurons follows daily pattern, are suppressed during feeding, and their inhibition acutely increases energy expenditure, body temperature, and activity levels. Long-term inhibition of this population produces sustained weight loss, selective reduction of white fat, preservation of lean mass and brown fat, and improved glucose tolerance even during high-fat feeding. These findings reveal a previously unrecognized circuit that links metabolic state with daily timing cues and suggest that targeting this neuronal population may offer new strategies for treating obesity and related metabolic disorders.

## Introduction

Energy homeostasis, balancing intake with expenditure, is critical for survival. Energy is taken in through feeding, a tightly regulated process that balances intake with caloric needs ^1–3^. Energy expenditure (EE) is also tightly regulated, but mechanisms controlling EE are less understood^4–6^. In addition to changes in EE due to physical activity and environmental temperatures, EE varies with circadian timing and in response to changes in caloric intake (e.g. diet-induced and adaptive thermogenesis)^7–12^. Governed by the central nervous system, energy balance is tightly controlled, but small imbalances can develop and compound over time to produce weight gain or loss. In a obesogenic environment, imbalances leads to obesity and metabolic disorders^13,14^. Modulation of EE is a long-sought therapeutic target for the treatment of obesity and associated metabolic disorders^15–18^. This is particularly true for therapies that may increase the potential health promoting function of brown adipose tissue (BAT) and mitigate adverse effects of excess white adipose tissue (WAT)^19–22^. Neural circuits regulating EE are a promising avenue for therapeutic development but remain poorly understood.

The preoptic area of the hypothalamus (POA) is a central hub for homeostatic functions and plays increasingly understood roles in energy homeostasis and metabolic regulation^23–28^. The POA integrates physiological signals and connects to multiple brain regions that drive coordinated physiological and behavioral responses. Recent studies have identified POA neural populations regulating thermogenic responses to environmental warmth or cold, driving febrile responses, and critical for the dramatic decreases in EE that occur as part of fasting-induced torpor in mice ^29–40^. Other studies have shown that modulation of activity of POA neurons can alter systemic glucose and fatty acid levels^41–44^. Metabolomic studies implicated opioid peptides and kappa opioid receptor (KOR) signaling in the POA in responding to sustained caloric restriction^39,45^. Additionally, KOR signaling in the CNS is a potent modulator of thermogenic energy expenditure^46,47^. Single nuclei RNA sequencing of POA cells indicates *Oprk1* (KOR) is expressed in a small subset of the many neural types in the POA.^48^

We sought to test possible roles of KOR expressing neurons (POA^KOR+^) in the POA in regulating energy balance. In the studies presented here, we quantified population calcium activity in POA^KOR+^ neurons using fiber photometry and found that POA^KOR+^ neurons were recruited during the light period of the day and were suppressed with the initiation of feeding, particularly during refeeding after food deprivation. We tested the functional roles of POA^KOR+^ neurons using chemogenetic tools and found that POA^KOR+^ neurons bidirectionally regulate EE, thermogenesis, and locomotor activity. Finally, chronic silencing of outputs of POA^KOR+^ neurons in chow fed mice led to sustained increases in EE without compensatory hyperphagia. The imbalance drove weight loss with selective reductions in white adipose tissue (WAT). Lean and BAT masses were comparatively preserved. Chronic silencing of POA^KOR+^ neurons in mice fed and maintained on a high fat diet (HFD) led to weight loss returning mice to pre HFD weights, reductions in adipose mass, improved glucose tolerance, and metabolic remodeling WAT. Together, these findings demonstrate that POA^KOR+^ neurons regulate energy balance, highlighting their potential as a therapeutic target to shift energy balance to treat obesity-associated metabolic disorders.

## Results

### POA^KOR+^ neuron activity follows a daily pattern

Prior studies have implicated KOR signaling and POA^KOR+^ neurons in both obesity and caloric restriction, and single nucleus RNA sequencing (snRNAseq) data on cells from POA indicated *Oprk1* is selective expressed by a narrow population of neurons ^47–50^. To discover what leads to recruitment of POA^KOR+^ neurons, we used fiber photometry in KOR-Cre mice. Adeno-associated viruses (AAVs) were used to express GCaMP7s in the POA, with optic fibers implanted to record calcium activity in POA^KOR+^ neurons (**Fig. 1b-c**). Fiber placements centered on lateral POA, and the GCaMP7s expression was limited to the POA region (**Extended Data Fig. 1a**). Along with recording Ca^2+^-dependent and isosbestic signals, we video recorded the animals, obtained high resolution feeding behavior using FED3 devices, and monitored wheel running activity for 24 hours. Food was freely available, as the FED3 device replaced each chow pellet automatically when taken (free feeding). Animals were housed in 12:12 light dark cycle. We quantified the frequency of Ca^2+^ events, using the conservative threshold of greater than two standard deviations (2SD) above baseline to minimize potential over and double counting. Ca^2+^ events occurred more frequently between 6:00 and 18:00, when the lights were on (light phase) than 18:00 and 6:00 when lights were off (dark phase) (**Fig. 1d-g**). Quantification of events per hour as a function of time of day revealed a peak in increase in the frequency of events during the middle of the light phase (**Fig. 1g**). Raleigh plots reveal a mean of Raleigh vector near 12:00, noon, for Ca^2+^ events (**Fig. 1h**, **i**). As expected, feeding (**Fig. 1j**) and wheel running were higher in the dark phase (**Fig. 1k**). Together these data demonstrate that POA^KOR+^ neurons are more active during the light phase, when both feeding and wheel running are lower.

**Fig. 1.**
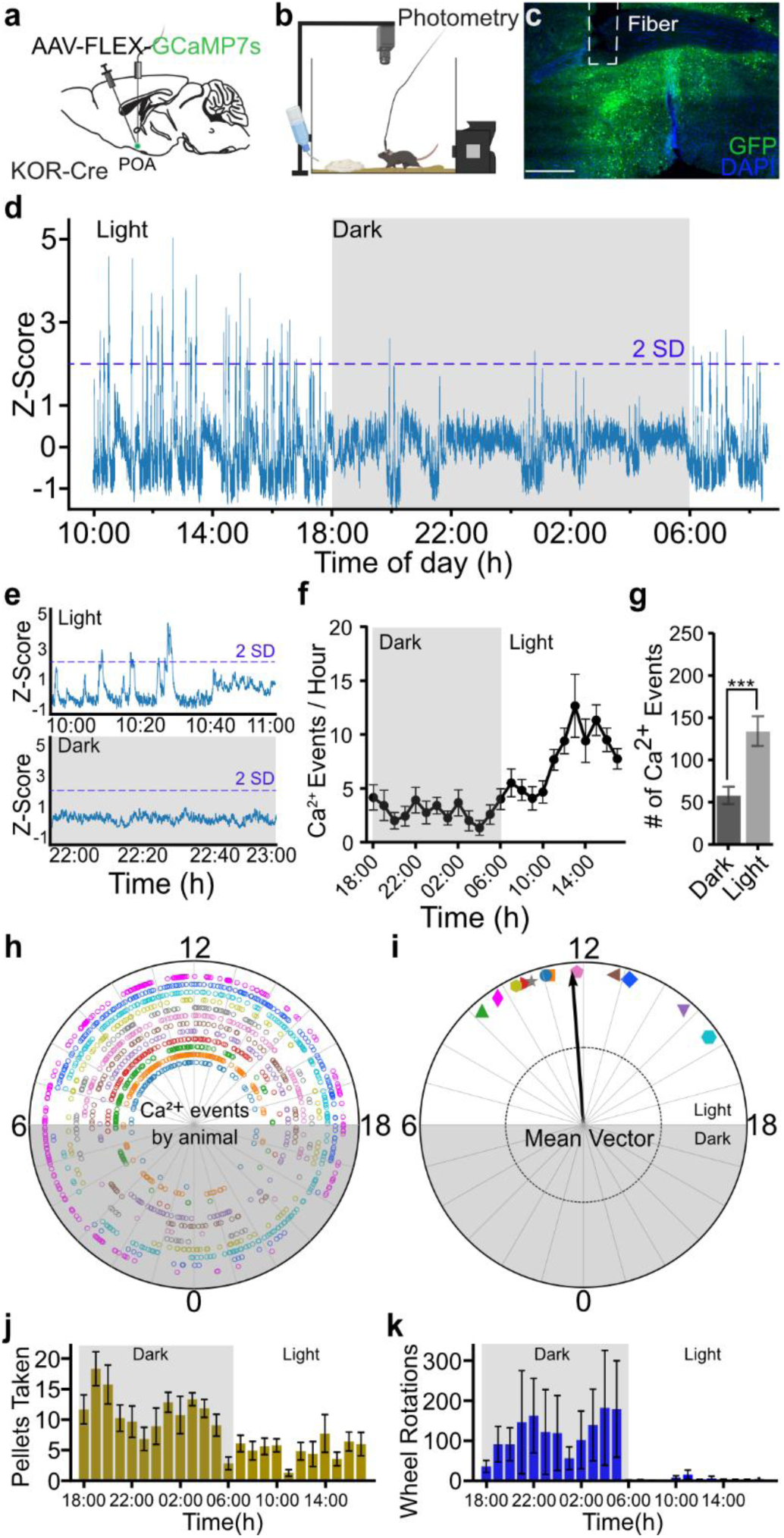
POA^KOR+^ neurons activity are active during the light phase. **a**, Schematic of virus injections of AAV-FLEX-GCaMP7s into the POA of KOR-Cre mice to label POA^KOR+^ neurons. **b**, Schematic of set-up for long-term fiber photometry and FED3 device recording. **c**, Representative images for POA^KOR+^ neurons infected with GCaMP7s GFP (green) and location of fiber placement. Scale bars, 250 μm. **d**, Representative original fiber-photometry recording of calcium events of POA^KOR+^ neuron under a 24-hour continuous recording condition. Z-score value was visualized 2 standard deviations (2SD) above baseline. **e**, Representative fiber-photometry recording of calcium events that occurred at light phase and dark phase. **f**, Distribution of calcium events per hour of POA^KOR+^ neuron under light and dark phase. **g**, Quantification of calcium events that occurred at light phase and dark phase. **h**, Raleigh plots of calcium events by individual animal. **i**, Mean Raleigh vector of calcium events distribution under light and dark phase. **j**, Mean food pellets taken per hour under light and dark phase. **k**, Wheel running activity recording under light and dark phase. Unpaired-sample t test in (**g**), Rayleigh test for non-uniformity of circular data and circular one-way ANOVA to compare circular data (n=12 mice). *P < 0.05, **P < 0.01, ***P < 0.001, and ****P < 0.0001; ns, no significance. Data are means ± s.e.m.

We monitored mice using FED3 devices to track feeding, video tracking to monitor movement, and running wheel activity. We found chow pellets were often taken in bouts with less than 30 seconds between pellets within a bout (**Extended Data Fig. 1b**). Using the 30 second inter pellet interval to define a bout, mice had approximately 40 feeding bouts in 24 hours. We found a small suppression of the Ca^2+^-dependent signal correlating with onset of a feeding bout (**Extended Data Fig. 1c**). We did not see any Ca^2+^-dependent signal changes at the onset of nest entrances or exits as movement controls, or at random time point as an additional control (**Extended Data Fig. 1d-f**).

### Ca^2+^ events in POA^KOR+^ neurons do not reflect the light state

Because mice were recorded in a 12:12 cycle, it is possible that the circadian like cycle of Ca^2+^ activity was entrained to the light cycle, and not an internal clock. To test this, we advanced the light cycle six hours while repeating the recordings. This did not alter the cycle of calcium signaling, nor total number of Ca^2+^ events, despite the earlier light on time on the advanced group (**Extended Data Fig. 2a-d**). Totals for pellets taken over 24 hours were not changed although the timing of feedings changes was altered by the light shift (**Extended Data Fig. 2e**, **f**). We conclude that Ca^2+^ event timing does not directly reflect the light state, which supports the possibility that recruitment of POA^KOR+^ neurons follows an internal circadian cue.

### Refeeding after food deprivation suppressed POA^KOR+^ neurons

Prior studies have shown increased activity of neurons in the POA in response to caloric limitation, notable during fasting induced torpor^23,32,33,51^. We therefore examined the impact of 18 hours of food deprivation followed by refeeding on POA^KOR+^ neuron activity (**Fig. 2a**, **b**). During food deprivation, there was no difference in frequency of Ca^2+^ events between the deprived and ad lib baseline condition (**Fig. 2c**, **d**). However, in the 5 hours after return of food there was marked suppression of Ca^2+^ events, which coincided with elevated feeding after the deprivation ended (**Fig. 2e**). This suggested that refeeding suppresses the activity of POA^KOR+^ neurons.

**Fig. 2.**
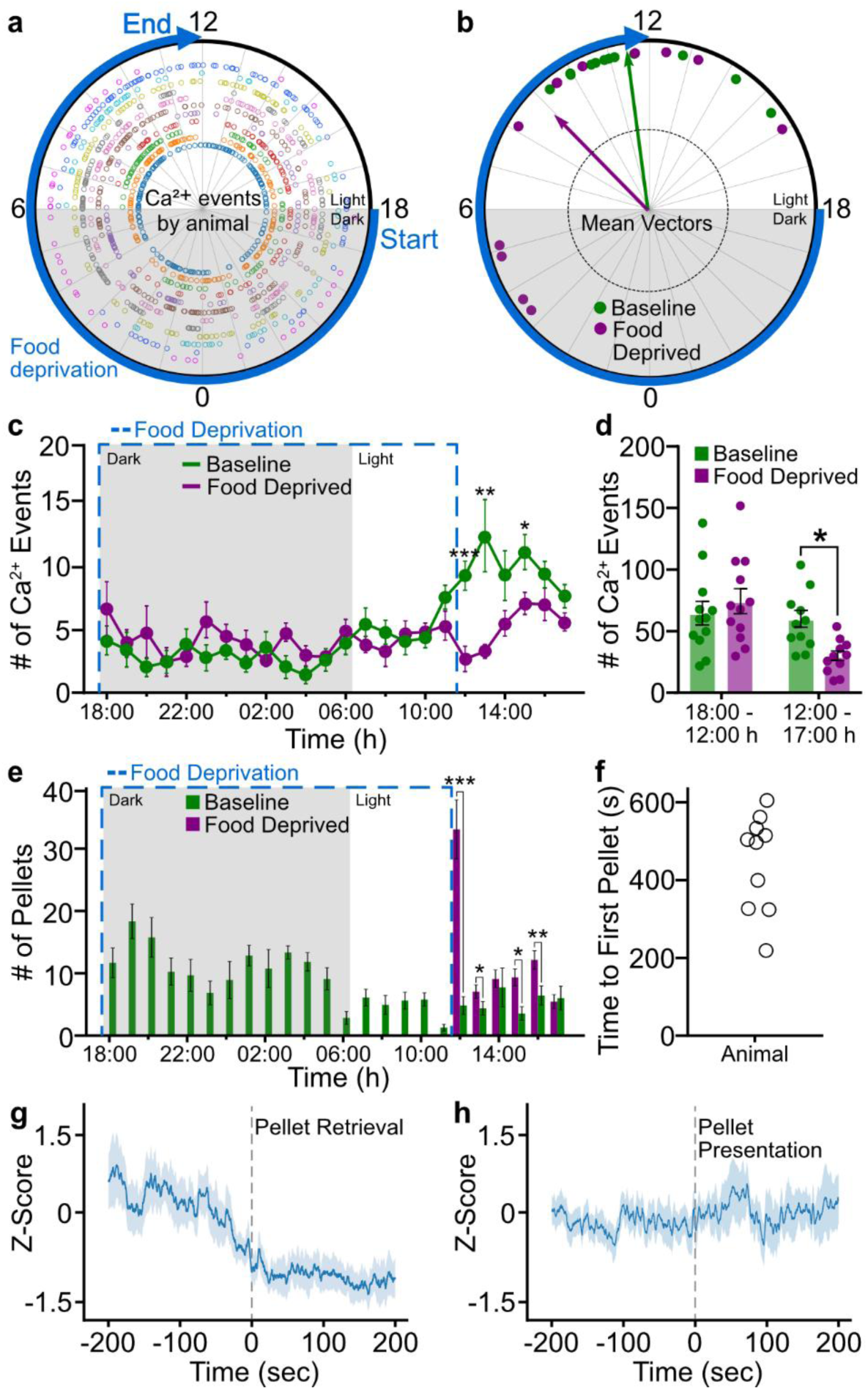
POA^KOR+^ neurons activity is suppressed by refeeding. **a**, Raleigh plots of calcium events by individual animal under an 18 hour fasting scheme. **b**, Mean Raleigh vector of calcium events distribution under an 18 hour fasting scheme. **c**, Distribution of calcium events per hour of POA^KOR+^ neuron under normal light-dark circle (Baseline) and fasting conditions. **d**, Comparison of number of calcium events recorded under normal light-dark cycle and fasting conditions. The time period from 18:00 till 12:00 the following day was for fasting and 12:00 till 17:00 the following day for refeeding. **e**, Comparison of number of food pellets taken per hour under normal light-dark cycle and fasting scheme. **f**, Time taken for retrieving the first pellet after fasting. **g**, Ca^2+^ signals of the first pellet retrieval after fasting. **h**, Ca^2+^ signals of pellets presentation after fasting. Rayleigh test for non-uniformity of circular data and circular one-way ANOVA to compare circular data (n=12 mice), two-way ANOVA comparing baseline and fasting events in (**c**), unpaired-sample *t* test in (**d**). *P < 0.05, **P < 0.01, ***P < 0.001, and ****P < 0.0001; ns, no significance. Data are means ± s.e.m.

After the restart of food availability, animals took variable amounts of time to remove the first pellet (**Fig. 2f**) allowing us to examine the temporal relationship of feeding behavior to POA^KOR+^ neuron recruitment. We aligned the Ca^2+^ dependent signal to the first pellet taken during refeeding and found suppression of the Ca^2+^ signal from POA^KOR+^ neurons (**Fig. 2g**). However, alignment of the Ca^2+^ -dependent signal to the availability of the first pellet in the FED3 device revealed no change in the Ca^2+^-dependent signal (**Fig. 2h**). Overall, we conclude that POA^KOR+^ neurons are suppressed by feeding, and a refeeding after fasting blocked the normal increases in Ca^2+^ signaling seen during the light portion of the day.

### POA^KOR+^ neuron recruitment not altered by changes in ambient temperatures

Neurons in the POA also respond to changes in ambient temperature, although snRNAseq data from POA suggest neurons expressing *Oprk1* are largely distinct^36,52–54^. We tested if POA^KOR+^ neuron responded to ambient warmth using Fos staining and fiber photometry (**Extended Data Fig. 3a, b**). KOR-Cre mice injected with AAV-DIO-eYFP to label POA^KOR+^ neurons were exposed to 38°C or room temperature (24°C) for 4 hours prior to harvesting brains. Immunohistochemistry revealed that exposure with 38°C increased Fos expression in the POA, but not in POA^KOR+^ neurons (**Extended Data Fig. 3c-e**). We also used KOR-Cre mice injected in the POA with AAV-Flex-GCaMP7s to examine change in recruitment of POA^KOR+^ neurons by warming or cooling the arena floor. Neither 34°C nor 10°C arena floor changed the Ca^2+^-dependent signal in POA^KOR+^ neurons (**Extended Data Fig. 3f, g**). This further indicates that POA^KOR+^ neurons are distinct from POA neurons that were implicated in responding to ambient temperature changes.

### POA^KOR+^ neurons express GABAergic markers

We next sought to understand the identity of POA^KOR+^ neurons. Using fluorescent in situ hybridization (FISH) we identified neurons in POA that labeled for *Oprk1* transcript and found 90% of these cells co-labeled for S*lc32a1* (VGAT) (**Extended Data Fig. 4a-d**). We made crosses of KOR-Cre mice to VGLUT2-Flp or VGAT-Flp mice and carried out combinatorial genetic experiments to further examine overlap of *Oprk1* expression with GABAergic or glutamatergic neurons. We injected AAV-DIO-mCherry mixed with AAV-Con/Fon-eYFP, which requires both Cre and Flp dependent recombination for expression of eYFP. In KOR-Cre x VGAT-Flp but not KOR-Cre x VGULT2-Flp mice we found robust expression of eYFP. In both cases Cre dependent expression of mCherry was present (**Extended Data Fig. 4e-l**).

### POA^KOR+^ neurons project to multiple brain regions linked with regulation of metabolism

To understand the anatomic relationships of POA^KOR+^ neurons, we examined the projections of POA^KOR+^ neurons by injecting AAV-DIO-eYFP in the POA, confirming selective labeling of cell bodies restricted to the POA, and imaging labeled processes (**Fig. 3a**, **b**). We identified projections from POA^KOR+^ neurons to many brain regions including multiple areas critical for regulating metabolism, which include the paraventricular nucleus of the hypothalamus (PVN), dorsal medial hypothalamus (DMH), periaqueductal grey (PAG), raphe pallidus (RPa), and parabrachial nucleus (PBN) (**Fig. 3c-h**). We further examined if POA^KOR+^ neurons had presynaptic terminals in key areas by injecting AAV-hSyn-FLEx-mGFP-2A-Synaptophysin-mRuby into the POA of KOR-Cre mice. After confirming the injections selectively targeted the POA, we identified for co-labeling by GFP and mRuby in bouton-like structures in areas including PVN, DMH, PAG, PBN, and RPa (**Extended Data Fig. 5a-f**). To test for possible anatomic subpopulations based selective projections from POA^KOR+^ neurons to different brain areas, we labeled POA^KOR+^ neurons (POA^KOR+^::DMH) projecting to the DMH, a key area for thermal and metabolic regulation, using retroAAV-DIO-Flp injected into the DMH and AAV-fDIO-eYFP injected into the POA of KOR-Cre mice. POA^KOR+^ labeled using with a combinatorial approach projected to the same brain areas as the total POA^KOR+^ population (**Extended Data Fig. 6a-h**) indicating POA^KOR+^ neuron arborize rather than have discrete subpopulations. Taken together, these data indicate that POA^KOR+^ neurons have arborizing projections to key brain areas implicated in regulating metabolism, feeding, and thermal homeostasis.

**Fig. 3.**
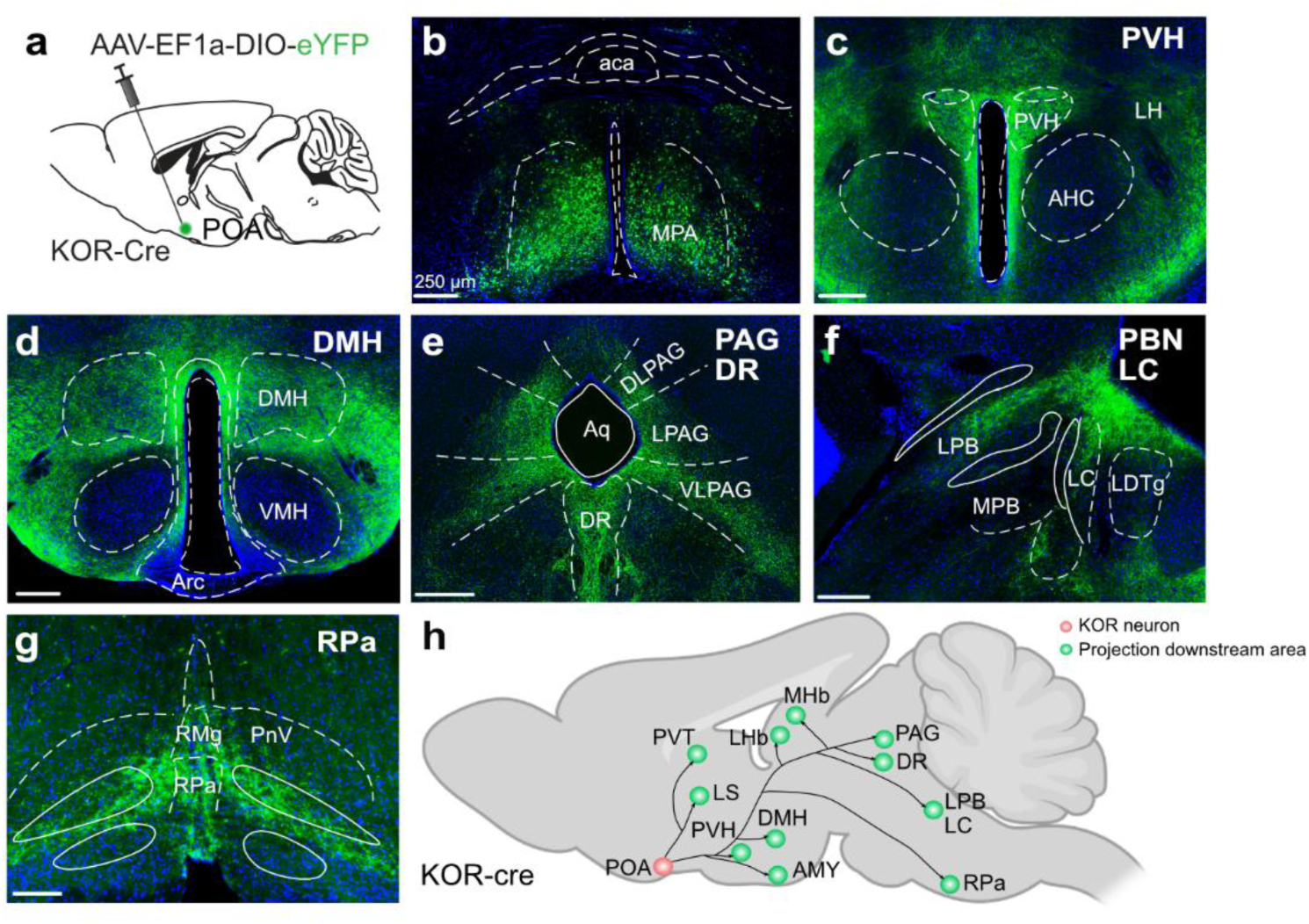
Whole-brain anterograde mapping of downstream metabolism regulatory circuits of KOR neurons. **a**, Schematic of virus injections of AAV-EF1a-DIO-eYFP into the POA of KOR-Cre mice to label POA^KOR+^ neurons. **b**, Cell bodies labeled with eYFP are limited to the medial side of POA. **c**-**g**, Distribution of downstream projections from POA^KOR+^ neurons. GFP-positive fibers are seen in multiple brain regions that are implicated in the regulation of body temperature and energy expenditure including the (**c**) PVH, (**d**) DMH, (**e**) PAG and DR, (**f**) PBN and LC, and (**g**) RPa. **h**, Summary of projection regions of POA^KOR+^ neurons. Major projection sites are indicated. AMY, amygdala; DMH, dorsomedial hypothalamus; DR, dorsal raphe; LC, locus coeruleus; LS, lateral septum; MHb, medial habenula; LHb, lateral habenula; PBN, parabrachial nucleus; PAG, periaqueductal grey; PVH, paraventricular hypothalamus; POA, preoptic area; PVT, paraventricular thalamic nucleus; RPa, raphe pallidus. Scale bars, 250 µm. n=4 mice.

### Inhibition of POA^KOR+^ neurons elevated energy expenditure

Finding projections from POA^KOR+^ neurons to multiple brain regions implicated in the regulating facets of metabolism and feed and the inhibition of POA^KOR+^ neurons correlating to feeding raises the possibility that POA^KOR+^ neurons regulate metabolic activity. To test the hypothesis that POA^KOR+^ neurons regulate metabolism, we expressed inhibitory Gi coupled Designer Receptors Exclusively Activated by Designer Drugs (Gi-DREADD) or eYFP in POA^KOR+^ neurons via injection of AAVs into the POA of KOR-Cre mice (**Fig. 4a**, **b**, and **Extended Data Fig. 7g**). Injections labeled KOR+ neurons in the POA with minimal labeling outside the POA. Mice with missed injections or labeling outside POA were excluded from analyses. Activation of inhibitory Gi-DREADDs by injection of CNO (1 mg/kg) led to increased temperatures of core, BAT, and eye (**Fig. 4c-f**), a validated surrogate for core body temperature^55^.

**Fig. 4.**
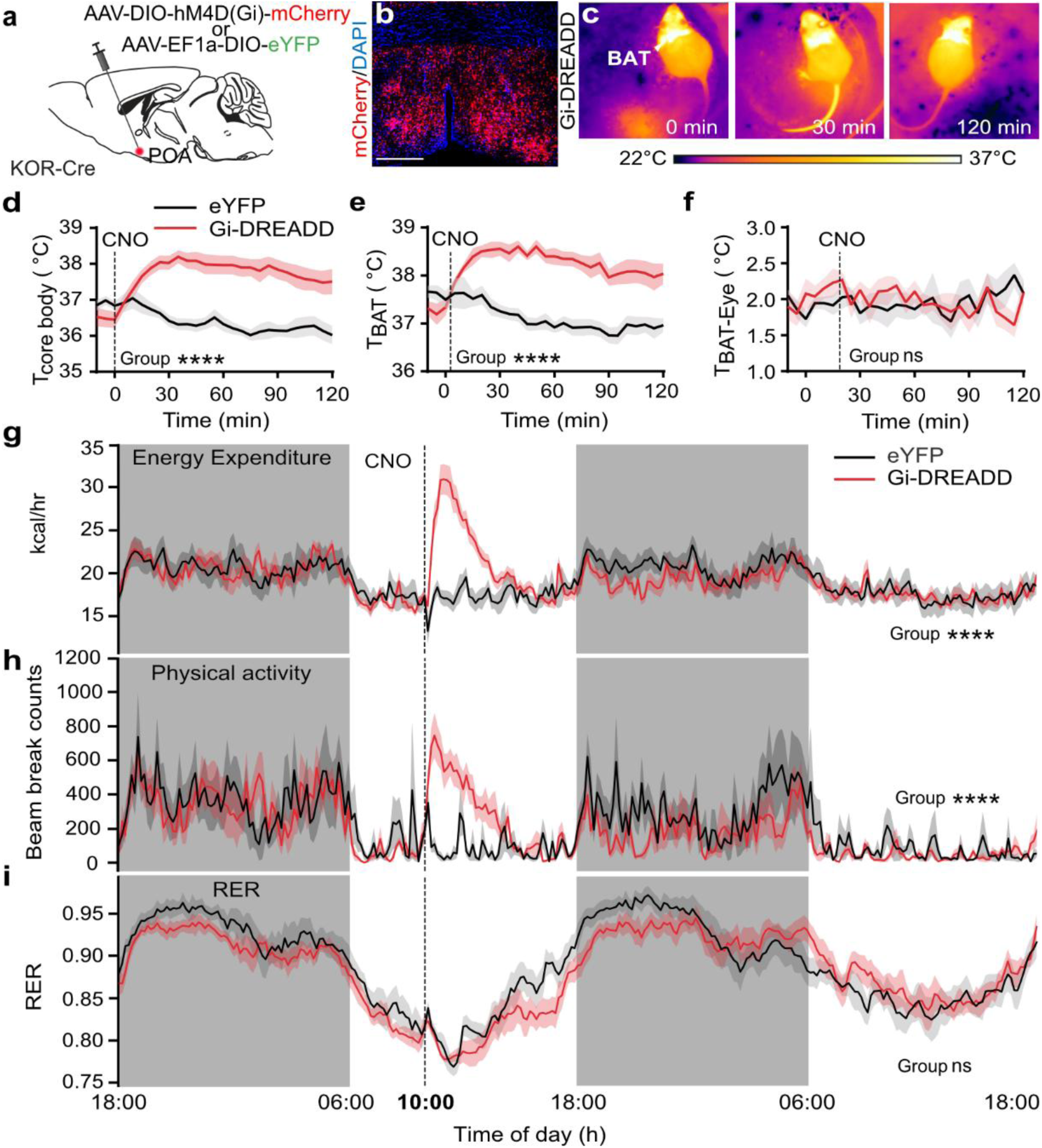
Chemogenetic inhibition of POA^KOR+^ neurons induces hypermetabolic rate and thermogenesis. **a**, Schematic of virus injections of AAV-hSyn-DIO-hM4D(Gi)-mCherry or AAV-EF1a-DIO-eYFP into the POA to inhibit POA^KOR+^ neurons. **b**, Representative images for POA^KOR+^ neurons infected with inhibitory DREADD hM4D (red). Scale bars, 250 μm. **c**, Representative thermal images of BAT before and after CNO (1 mg/kg, i.p.) injection. **d**, **e**, Chemogenetic inhibition of POA^KOR+^ neurons increases both core body temperature (**d**) and BAT temperature (**e**). **f**, Difference between BAT and eye temperatures showing largely overlapping indicating that BAT contributes to the increased body temperature. **g-i**, Chemogenetic inhibition of POA^KOR+^ neurons increases EE (**g**) and locomotor activity (**h**), not RER (**i**). CNO (1 mg/kg, i.p.) injection at 10:00 am in the morning. Two-way ANOVA comparing control and treated groups after CNO injection (n=7-10 mice per group). *P < 0.05, **P < 0.01, ***P < 0.001, and ****P < 0.0001; ns, no significance. Data are means ± s.e.m.

Prior studies have found that activation of temperature responsive POA neurons altered thermal preferences of mice. Artificial activation of warm-activated neurons drives lower body temperature and, paradoxically, preference for cooler temperatures^37^. In thermal gradient preference tests (**Extended Data Fig. 7d-f**), inhibition of POA^KOR+^ neurons shifted the thermal preference to much colder zones. Mice spent more time in the coldest zones (approximately 12°C) and less time in the warmer zones, around 30°C, compared to control mice. These data suggest that mice seek to correct for the elevation of body temperatures induced by acute inhibition of POA^KOR+^ through behavioral thermal homeostatic behaviors and do not support a primary role for POA^KOR+^ neurons in thermal regulation^36^.

To examine metabolic changes induced by inhibition of POA^KOR+^ neurons, we used metabolic profiling, focused first on 10:00 when POA^KOR+^ neurons were most active (**Fig. 1f**). Following injections of CNO, energy expenditure (EE) increased in Gi-DREADD expressing mice compared to fluorophore control mice (**Fig. 4g-i**). Locomotor activity was also increased. Respiratory exchange ratio (RER) was similar in mice from both conditions; however, this time of day corresponds to a normal circadian nadir in the RER, and detection of changes may be limited by a floor effect. We tested the effects of POA^KOR+^ neuron inhibition at a second time of day using separate cohorts of mice. CNO was injected at 18:00 (**Extended Data Fig. 7a-c**). Both EE and locomotor activity increased following CNO injection. The RER decreased, suggesting greater utilization of lipids for oxidation compared to controls. These data indicate POA^KOR+^ neurons can regulate EE through thermogenesis and locomotion.

### Activation of POA^KOR+^ neurons suppressed energy expenditure

In KOR-Cre mice we expressed excitatory Gq coupled DREADDs (Gq-DREADDs) or fluorophore control in POA^KOR+^ neurons using AAV injections in the POA (**Fig. 5a**, **b** and **Extended Data Fig. 8g**). POA^KOR+^ neuron activation induced by CNO (0.5 mg/kg) induced rapid decreases in body and BAT temperatures in Gq-DREADD expressing mice compared to control (**Fig. 5c-e**). To examine the duration of the hypothermia further, we implanted mice with temperature logging devices and found long lasting suppression of body temperature taking 32 hours after CNO administration to fully recover (**Fig. 5f**).

**Fig. 5.**
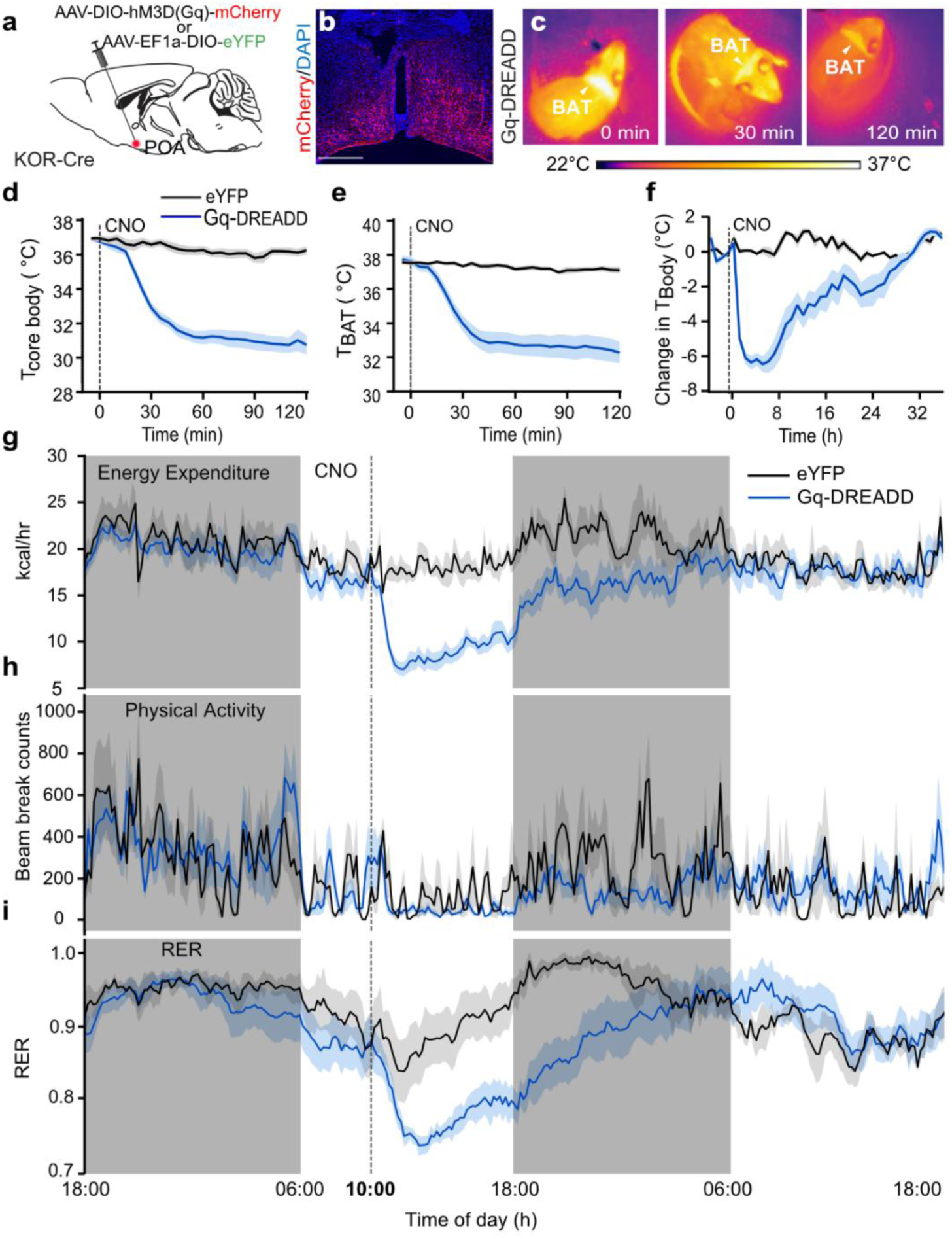
Chemogenetic activation of POA^KOR+^ neurons induces hypometabolic state and hypothermia. **a**, Schematic of virus injections of AAV-hSyn-DIO-hM3D(Gq)-mCherry or AAV-EF1a-DIO-eYFP into the POA to activate POA^KOR+^ neurons. **b**, Representative images for POA^KOR+^ neurons infected with excitatory DREADD hM3D (red). Scale bars, 250 μm. **c**, Representative thermal images of BAT before and after CNO (0.5 mg/kg, i.p.) injection. **d**, **e**, Chemogenetic activation of POA^KOR+^ neurons suppress both core body temperature (**d**) and BAT temperature (**e**). **f**, Long-term changes of core body temperature monitored with iButton implanted mice. **g-i**, Chemogenetic activation of POA^KOR+^ neurons suppress EE (**g**) and locomotor activity (**h**), and RER (**i**). CNO (0.5 mg/kg, i.p.) injection at 10:00 amTwo-way ANOVA comparing control and treated groups after CNO injection (n=7-12 mice per group). *P < 0.05, **P < 0.01, ***P < 0.001, and ****P < 0.0001; ns, no significance. Data are means ± s.e.m.

We examine the effects of POA^KOR+^ neuron activation on metabolic activity in a separate cohort of mice. Activation of POA^KOR+^ neurons at 10:00 by CNO (0.5 mg/kg) injection caused EE to decline markedly in Gq-DREADD expressing mice compared to control **(Fig. 5g**). Suppression of EE was long lasting. During the light phase control mice had low overall locomotor activity; however, there was a decline in locomotor activity in the Gq-DREADD mice. In the subsequent dark phase, when control mice became more active, locomotor activity in the Gq-DREADD expressing mice was lower (**Fig. 5h**). RER was lower following activation of POA^KOR+^ neurons and returned to control values approximately 20 hours after injection of CNO (**Fig. 5i**). In a separate cohort of mice, we injected CNO at 18:00, a time of normally low Ca^2+^ event frequency, and found similar change, but all measures returned to baseline more quickly, less than 12 hours after CNO injection (**Extended Data Fig. 8a-c**). Activation of POA^KOR+^ neurons also led to a small shift to cooler thermal preference and a decrease in the total distance moved (**Extended Data Fig. 8d-f**). Together, these data indicate that activation of POA^KOR+^ neurons leads to a hypometabolic state and an increase in use of lipids as a metabolic substrate.

### Silencing of POA^KOR+^ neurons drove weight loss and increased energy expenditure

Increasing energy expenditure has been a long-sought possible therapeutic methodology for weight loss and metabolic dysfunction. Our data indicated that inhibiting POA^KOR+^ neurons can increase EE, although the effect was transient. To test if chronic inhibition of POA^KOR+^ neuronal output would cause sustained reductions in metabolic activity, we disrupted synaptic release from POA^KOR+^ neurons by expressing tetanus toxin light chain (TetTox) in POA^KOR+^ neuron using AAV injections into the POA of KOR-Cre mice (**Fig. 6a** and **Extended Data Fig. 9e**). We examined feeding data using FED3 devices set to a fixed ratio of 1 (FR1), such that mice received one 20 mg chow pellet each time they poked in the nose port. Mice were trained prior to viral injections (**Fig. 6b**). After viral injections TetTox expressing mice started losing weight compared to eYFP control mice (**Fig. 6c**, **d**). Weight loss plateaued nine days after viral injections. Body composition analyses demonstrated decreased fat mass. Loss of lean mass did not reach significance for all mice together (**Fig. 6e**, **f**). However, variance in lean mass was increased because of differences between male and female mice. Analyses by sex revealed a significant weight loss and lower fat mass in male and female mice. A lower lean mass was significant for male and female mice when analyzed by sex (**Extended Data Fig. 10a-h**). Among major adipose depots, inguinal (iWAT) and epididymal adipose tissues (eWAT) were significantly smaller in TetTox expressing mice, but the BAT mass was preserved **(Fig. 6g-i**). Analyses of cumulative and daily average pellet retrieval revealed that TetTox expressing mice did not take significantly fewer pellets than control mice (**Fig. 6j**, **k**), if anything they ate slightly more pellets, although not significant. We repeated these studies using independent cohorts with FED3 devices set to free feed, which automatically replaced a pellet when one was taken without port activation required (**Extended Data Fig. 11a-f**). As in the FR1 cohort, under free feeding conditions TetTox expressing mice lost weight while control mice did not (**Extended Data Fig. 11a, b**). Despite the weight loss, under free feed conditions pellet retrieval was not significantly different between the TetTox and control groups (**Extended Data Fig. 11c, d**). These data indicate that silencing POA^KOR+^ neurons drives weight loss, but the weight lost is not due to reductions in feeding.

**Fig. 6.**
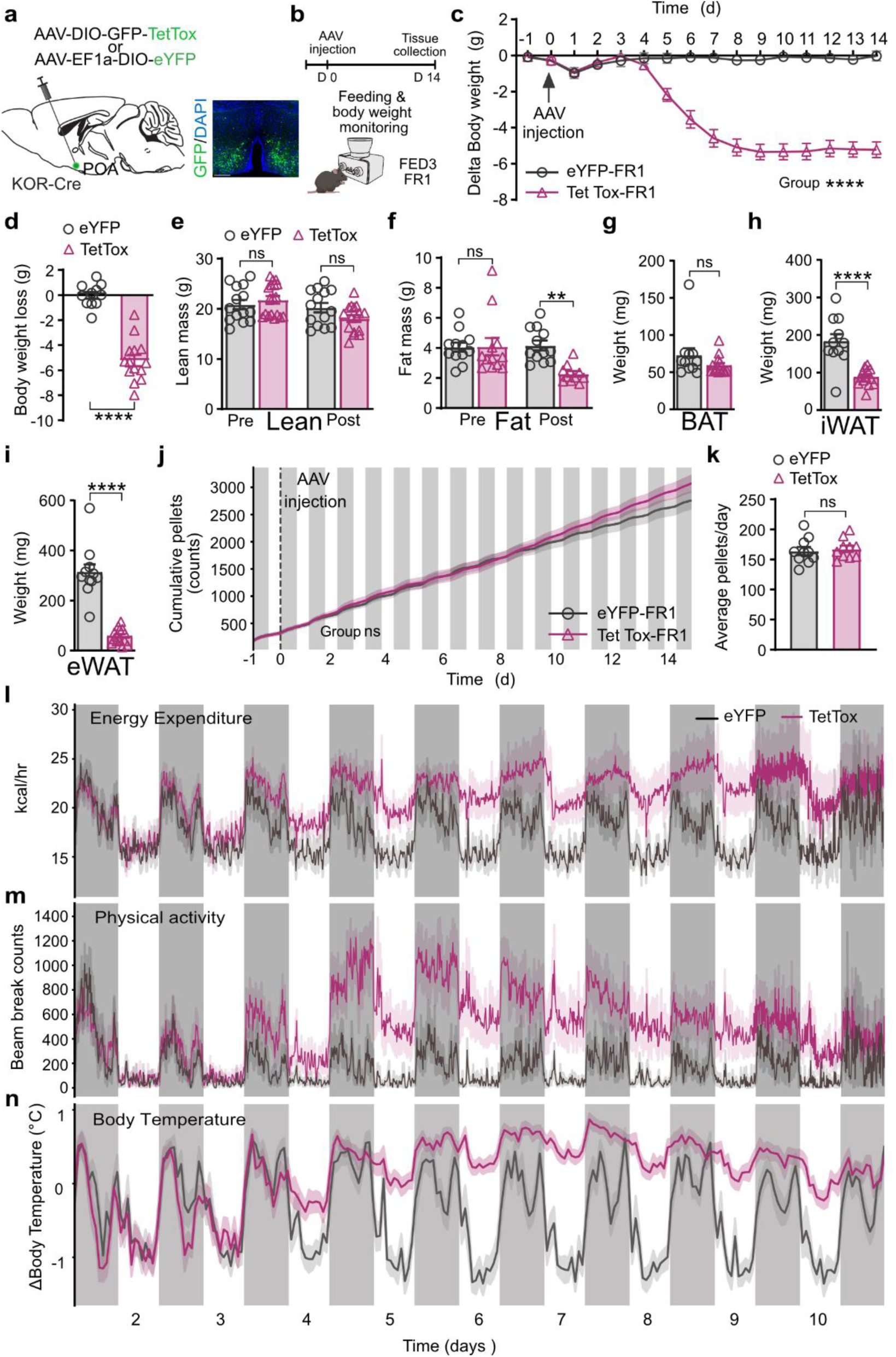
Synaptic blockade of POA^KOR+^ neurons promoted energy expenditure and causes body weight loss. **a**, Schematic of virus injections of AAV-DIO-GFP-TetTox or AAV-EF1a-DIO-eYFP into the POA to chronically block synapse transmission of KOR neurons. Representative images showing the expression of TetTox (green) in POA^KOR+^ neurons (right). Scale bars, 250 μm. **b**, Experimental timeline for virus injection, body weight measurement, feeding monitored with FED3 device, and tissue collection after silencing KOR neurons in the POA. **c**, Body weight changes after silencing POA^KOR+^ neurons showing mice start to lose body weight 5 days after TetTox injection and enter a steady state after 10 days. **d**, Average body weight changes between control and TetTox treated animals. **e**, **f**, Fat (**e**) and lean (**f**) masses of control eYFP or TetTox treated mice were measured by EchoMRI at two weeks after viral injection. **g-i**, Quantification of tissue organ weight of BAT (**g**), iWAT (**h**), and eWAT (**i**) after silencing KOR neurons. **j**, Cumulative food intake monitored with FED3 device between control and TetTox treated mice over the two weeks recording period. **k**, Average food pellets taken per day between control and TetTox treated animals. **l**, Energy expenditure showed constant increase after silencing POA^KOR+^ neurons. **m**, Physical activity showed increase at early timepoint and then began to drop after silencing POA^KOR+^ neurons. **n**, Body temperature changes monitored with iButton implantation showing elevated body temperature after viral injection. Two-way ANOVA comparing control and treated groups in (**c**), (**j**), (**l**), (**m**), and (**n**). Unpaired-sample *t* test in (**d-i**) and (**k**). *P < 0.05, **P < 0.01, ***P < 0.001, and ****P < 0.0001; ns, no significance. n=10-14 mice per group. Data are means ± s.e.m.

Metabolic phenotyping measurements demonstrated that silencing POA^KOR+^ neurons led to increased EE. Elevations of EE were first evident during the dark phase of post injection day three and continued through post injection day 10 (**Fig. 6l** and **Extended Data Fig. 9f-h**). The difference in EE was most prominent during the light phase when EE declined in control but not TetTox mice. Quantification of physical activity revealed increased activity in TetTox mice starting post injection day 3 (**Fig. 6m**). Physical activity was elevated during both the light and dark phases. Unlike EE however, physical activity began returning to normal ranges on days nine and 10. The RER was lower starting five days after viral injections, likely reflecting use of fat stores because EE outstripped intake (**Extended Data Fig. 9a**). RER normalization coincided with weight stabilization. Data from implanted temperature logging devices in separate KOR-Cre mice revealed TetTox expressing mice had altered body temperatures day four after viral injection (**Fig. 6n**). Body temperatures were elevated compared to controls during the light phase of the day when body temperatures declined in control mice because the daily decline in body temperature was blunted in TetTox expressing mice. TetTox and control mice had similar thermal preference during the light phase (**Extended Data Fig. 9b-d).** The data from feeding, EE, and body temperature recordings indicated that silencing POA^KOR+^ neurons leads to weight loss because of increased EE due to elevated thermogenic and physical activity.

### Silencing POA^KOR+^ neurons drove weight loss in mice with high fat diet induced obesity maintained on high fat diet

To test if silencing POA^KOR+^ neurons would alter weight in mice fed a high fat diet (HFD), KOR-Cre mice were maintained on HFD for 10 weeks prior to viral injections. 10 weeks of HFD induced weight gain in both male and female mice (**Extended Data Fig. 12b**). Males gained more weight (**Extended Data Fig. 12c**) and because of the differences in weight gain following HFD male and female mice were analyzed separately. Mice were maintained on HFD after viral injection of AAVs to express TetTox or eYFP into the POA of KOR-Cre mice (**Fig. 7a**, **b**). Mice expressing TetTox lost weight while control mice gained weight over the three-weeks following viral injections (**Fig. 7c** and **Extended Data Fig. 13a**). The weights of the two groups diverged five days after viral injections and stabilized during the 21 days of measurement. Both male and female TetTox mice returned to lean pre-HFD weights despite the ongoing HFD, while control mice became heavier over the same 21-day period (**Extended Data Fig. 12f, g** and **Extended Data Fig. 13m**, **n**). Food intake was monitored daily. TetTox expressing HFD fed mice had a transient decrease in feeding with maximum suppression occurring on day 6; but feeding returned to same levels as controls (**Fig. 7d** and **Extended Data Fig. 13b**). Despite the return of baseline feeding matching controls mice, TetTox expressing mice continued to lose weight. Activity measured using home cage monitoring revealed greater percentage of time active in TetTox expressing mice (**Extended Data Fig. 12a** and **Extended Data Fig. 13c**) during both the light and dark phases. Silencing POA^KOR+^ neurons in the context of excess caloric storage led to decreased transient feeding, in contrast to the data obtained using lean mice.

**Fig. 7.**
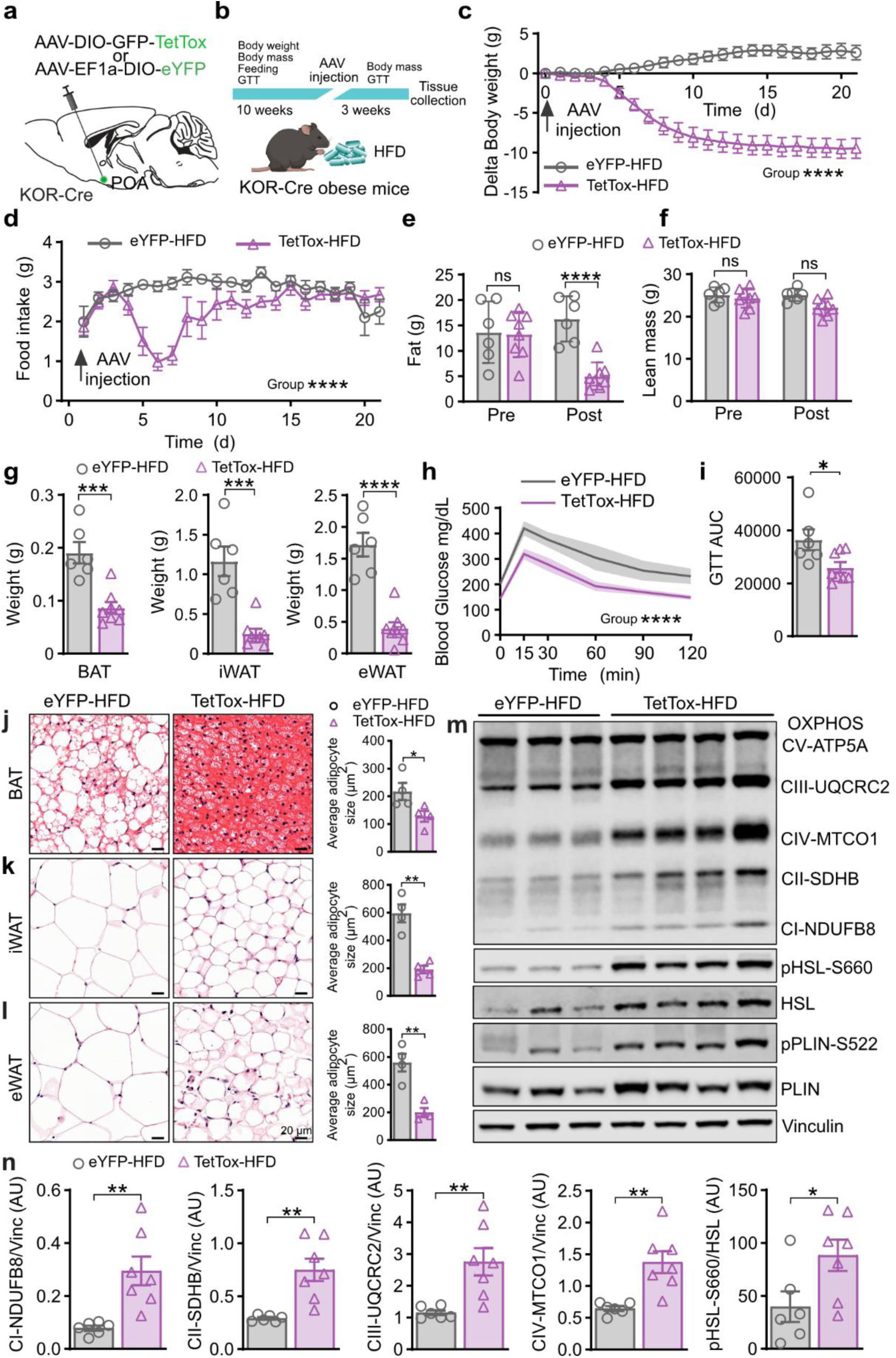
Synaptic blockade of POA^KOR+^ neurons prevented obesity and improves metabolic health. **a**, Schematic of virus injections of AAV-DIO-GFP-TetTox or AAV-EF1a-DIO-eYFP into the POA to chronically block synapse transmission of KOR neurons in obese mice. **b**, Experimental timeline for virus injection, body weight measurement, GTT, feeding, and tissue collection after silencing KOR neurons in the obese mice. **c**, Body weight changes after silencing POA^KOR+^ neurons showing mice start to lose body weight 5 days after TetTox injection and enter a steady state after two weeks. **d**, Average food intake per day between control and TetTox treated obese mice. **e**, **f**, Fat (**e**) and lean (**f**) masses of control eYFP or TetTox treated obese mice were measured by EchoMRI at two weeks after viral injection. **g**, Quantification of tissue organ weight of BAT, iWAT, and eWAT after silencing KOR neurons at three weeks after viral injection. **h**, Time course of blood glucose levels during the GTT in control and TetTox treated mice. **i**, Quantification of the AUC of GTT. **j**-**l**, H&E staining and quantified adipocyte size of BAT (**j**), inguinal WAT (iWAT) (**k**), and epididymal WAT (eWAT) (**l**), of control and TetTox treated KOR-Cre mice. **m**, Western blot of OXPHOS, HSL, and PLIN1 proteins in iWAT depots of control and TetTox treated mice. **n**, The quantified ratio of OXPHOS, pHSL, and PLIN1 for both groups. Statistical analyses included Two-way ANOVA comparing control and treated groups in (**c**), (**d**) and (**h**). Unpaired-sample *t* test in (**e**), (**f**), (**g**), (**i**), (**j**-**l**) and (**n**). *P < 0.05, **P < 0.01, ***P < 0.001, and ****P < 0.0001; ns, no significance. n=6-8 mice per group. Data are means ± s.e.m.

### Silencing POA^KOR+^ neurons led to improved glucose tolerance

Glucose tolerance was impaired in both male and female mice following 10 weeks of HFD (**Extended Data Fig. 12d, e** and **Extended Data Fig. 13o**, **p**), compared to age matched lean controls. Expression of TetTox in POA^KOR+^ neurons lead to improved glucose tolerance compared to eYFP controls in males (**Fig. 7h**, **i**) and in females (**Extended Data Fig. 13g, h**) or compared to previral injection. No improvement in glucose tolerance was evident in control mice (**Extended Data Fig. 12h-k** and **Extended Data Fig. 13i-l**). Although silencing POA^KOR+^ neurons improved glucose tolerance, further studies are required to understand the extent to which this is a direct effect of change in neural activity, direct changes in metabolism, or secondary to weight loss.

### Silencing POA^KOR+^ neurons led to remodeling of adipose tissues while on HFD

Following viral injections TetTox expressing mice lost significantly more fat mass than controls. Lean mass was preserved (**Fig. 7e**, **f** and **Extended Data Fig. 13d, e**). Measurements of specific adipose tissue depots revealed the masses of BAT, eWAT, and iWAT tissues decreased in TetTox but not control mice (**Fig. 7g** and **Extended Data Fig. 13f**). Histological examination demonstrated that on HFD BAT developed unilocular droplets and increased in size in control mice (**Fig. 7j**). Silencing POA^KOR+^ neurons led to reversal of these morphologic changes and decreased adipocyte size. Adipocyte size also decreased in eWAT and iWAT tissues (**Fig. 7k**, **l**). Biochemical data from iWAT tissues revealed that the components of the mitochondrial electron transport chain, ubiquinol–cytochrome c reductase core protein II (UQCRC2), mitochondrially encoded cytochrome C oxidase I (MTCO1), succinate dehydrogenase (SDH), NADH dehydrogenase ubiquinone oxidoreductase subunit B8 (NDUFB8) were significantly increased in mice expressing TetTox in POA^KOR+^ neurons (**Fig. 7m**, **n**). Changes in ATP synthase F1 subunit alpha (ATP5A) did not rise to the level of significance (**Extended Data Fig. 12l**). Additionally, the level of phosphorylated (active) hormone sensitive lipase (HSL) was significantly increased, suggesting an increase in adipose tissue lipolysis. In conclusion, silencing of POA^KOR+^ neurons led to loss of adiposity and WAT metabolic remodeling consistent with greater metabolic activity and lipolysis in adipocytes.

## Discussion

Sustained positive energy balance leads to obesity and related metabolic disorders. Once gained, the increased body mass is then defended both by feeding and metabolic adaptations that lower EE^56–61^. Identifying and targeting neural circuits able to increase EE, particularly without triggering compensatory food intake, would be of tremendous therapeutic value. Our studies sought to identify neural pathways that function in the regulation of EE and were focused on POA^KOR+^ neurons due to the evidence indicating KOR signaling and potentially POA^KOR+^ neurons can alter EE^46,49,62^.

The results presented here identify POA^KOR+^ neurons as a GABAergic population (**Extended Data Fig. 4**). Whole-brain mapping demonstrated arborizing projections from POA^KOR+^ neurons to multiple brain areas (**Fig. 3** and **Extended Data Figs. 5** and **6**) including the PVN, DMH, PAG, PBN, and RPa, key nodes in circuits regulating feeding, thermogenesis, EE, and metabolism ^31,32,63–65^. We found that POA^KOR+^ neurons were recruited following a temporal pattern that peaked during the light phase, the rest period for mice (**Fig. 1**). However, acute shift of the light schedule had only minor effects on the recruitment of POA^KOR+^ neurons, indicating light is not the primary driver (**Extended Data Fig. 2**). During food deprivation POA^KOR+^ neurons did not display an increased frequency of Ca^2+^ events but were suppressed immediately prior to refeeding, but suppression did not correspond to food presentation (**Fig. 2**). These data suggest that POA^KOR+^ neurons respond to internal cues related to time-of-day anticipation of feeding rather than detection of caloric intake (**Fig. 2** and **Extended Data Figs. 1** and **2**).

Activation or inhibition of POA^KOR+^ neurons yielded bidirectional changes in EE, locomotor activity, and body temperatures (**Figs. 4** and **5**, and **Extended Data Figs. 7** and **8**). DREADD mediated inhibition of POA^KOR+^ neurons increased EE at both 10:00 and at 18:00 indicating tonic regulation by POA^KOR+^ neurons. Supporting a functional link to circadian timing, the length of the and magnitude of effects of POA^KOR+^ interacted with the time-of-day activated (**Fig. 5** and **Extended Data Fig. 8**). Chronic silencing of POA^KOR+^ neurons led to weight loss in both lean mice and mice with HFD induced obesity. In lean mice a sustained increase in EE was evident with mild elevation in body temperatures, which was most evident during the light phase, leading to blunting of daily changes in body temperature (**Fig. 6**). Other means of elevating EE like cold ambient temperatures or induced thermogenesis are followed by hyperphagia^46,66–68^. Although EE was elevated following silencing of POA^KOR+^ neurons, feeding over the three weeks we examined was not changed in lean mice while body weight stabilized at a lower level. In mice fed HFD chronic silencing of POA^KOR+^ neurons drove weight loss, preserved lean mass, decreased WAT mass and metabolic remodeling of WAT depots (**Fig. 7j-n**). Understanding the underlying reasons for the transient decrease in food intake in HFD fed mice will require further studies. Because WAT shows signs of increase lipolysis, an avenue of future studies will be to determine if high levels of energy release from adipose stores in HFD fed mice leads to transient suppression of feeding. Together the presented data demonstrate POA^KOR+^ neurons are active under basal conditions, can regulate EE, are likely linked to circadian rhythms, and can promote EE without triggering compensatory feeding. Chronic modulation can promote weight loss through selective reduction of WAT. Thus, modulation of POA^KOR+^ neurons may be a viable target to improve metabolic health.

Excessive adiposity drives metabolic syndrome, increases inflammation, causes organ dysfunction, and can increase cancer risk ^82–87^. Although the role of BAT in humans remain an active area of research, increased BAT or metabolically active adipose tissue is associated with better metabolic health and may play a causative role in this association^88–92^. Silencing POA^KOR+^ neurons led to weight loss with preserved lean body mass in lean and obese mice. In lean mice, BAT mass was preserved, and WAT was markedly decreased. In mice on HFD, silencing POA^KOR+^ restored BAT towards a pre HFD state with reduced adipocyte, decreased lipid accumulation, and preservation of multilocular lipid droplet morphology. WAT showed increased potential for metabolic activity and evidence of elevated lipolysis. The selective diminution of WAT, even while maintained on HFD, highlights the potential positive impact of modulating the involved pathways. However, further studies are required to elucidate the underlying mechanisms and neural circuits responsible for these WAT-specific effects. Notably, the observation that chronic modulation of POA^KOR+^ neurons increased energy expenditure without concomitant increases in food intake provides a framework for future investigations aimed at overcoming maladaptive defenses that maintain elevated body mass.

The preoptic region of the hypothalamus is a major integratory hub, and recent studies have described warm responsive neurons defined by expression of *Adcyap1*, *Bdnf*, and *Lepr* that, when activated, lower body temperature^23,37,69^. Available data indicate POA^KOR+^ neurons are distinct from these populations. Warm responsive neural populations are largely glutamatergic. In contrast, we found POA^KOR+^ neurons express GABAergic markers and were not responsive to changes in environmental temperature. A prior study found that activation of POA GABAergic neurons located in the ventral lateral area (vLPO) can cause hypothermia and that GABAergic neurons in the POA projecting to the DMH were recruited by cold^70^. We did not find any data indicating POA^KOR+^ neurons respond to the thermal challenges (**Extended Data Fig. 3**). The long-lasting hypothermic and hypometabolic state induced following DREADD activation of POA^KOR+^ neuron resembled the effects of neural populations linked to torpor and torpor-like states marked by expression of markers including *Qrfp*, *Adcyap1,* and *Esr1*^23,32,33,40,51^. These glutamatergic populations are recruited torpor bouts brought on by food deprivation. Expression of the prostaglandin EP3 receptor (*Ptger3*), linked to fever production, is a unifying marker and *Ptger3* expressing neurons can function as a thermoregulatory switch capable of driving torpor and fever^38,40^. They also express *Slc17a6* (VGLUT2), which did not overlap with *Oprk1* in the studies here. Inhibition of POA^KOR+^ neurons raised body temperatures, as seen in febrile states, but mice also became hyperactive and sought markedly colder temperatures (**Extended Data Fig. 7**). The data weigh against linking POA^KOR+^ neurons to a state of febrile malaise, because fever and related circuits promote elevated thermal preference and lower activity^71^. An additional relevant population in the POA is the *Brs3* expressing group. They are a mixed GABAergic and glutamatergic population. Notably, when activated they have a biphasic effect with an initial decline in body temperature followed by an elevation in body temperature^30^. Published studies on *Brs3* expressing neurons focused on responses to cold and the induction of hyperthermia. Single nucleus sequencing data from the POA suggest the *Oprk1* expression partially overlaps with *Brs3* expression and there could be a commonality particularly among neurons that drive the decline in body temperature^48^. EE and metabolism are very closely linked to body temperature with maintenance of body temperature often representing a third of EE at typical room temperatures ^7,72^. Thermogenesis from metabolic processes contributes to body temperature and modulation of metabolic activity, not directly linked to thermoregulation, can impact body temperature. In the opposite direction, increases or decreased thermogenesis change EE and metabolic activity^4^. In context of knowledge of other POA populations, the results presented here show that POA^KOR+^ neurons are distinct GABAergic population that has daily patterns in recruitment and profound effects on EE. Thus, POA^KOR+^ neurons represent a node for regulating EE distinct from previously identified nodes.

Circadian timing exerts a fundamental influence on metabolism, synchronizing metabolic processes with daily time, environmental cues, and behavioral cycles^73,74^. This synchronization anticipates metabolic demands and adjusts fuel storage and mobilization optimizes metabolism during the active phase and help mitigate energy deficits during the rest phase^75,76^. Disruption of circadian synchrony, for example by shift work, which misaligns feeding, activity and intrinsic circadian timing, can worsen metabolic health^77–79^. Here we found that POA^KOR+^ neuron recruitment peaked during the rest phase and activation lowered EE, indicating metabolic changes (**Figs. 1** and **5**). Chronic silencing of POA^KOR+^ neurons damped daily rhythms in EE and body temperatures. POA^KOR+^ neurons were suppressed by feeding. Taken together, these findings suggest that POA^KOR+^ neurons may represent a previously unrecognized link between circadian timing, feeding time, and metabolic health.

Collectively, the data presented support a model in which POA^KOR+^ neurons function as a circadian-tuned regulator of energy balance, integrating internal metabolic signals with daily timing cues to shape thermogenesis, activity, and substrate utilization. Their distinct profile and anatomical connectivity set them apart from canonical warm-responsive or torpor-associated neurons, pointing instead to a specialized role in controlling basal EE. Modulation of this population meaningfully influences adipose tissue deposition. By demonstrating that long-term modulation of this population can alter body composition and metabolic health even under obesogenic conditions, our work highlights the broader principle that discrete hypothalamic nodes can recalibrate defended energy stores.

## Methods

### Mice

All animals were maintained, handled, and processed in accordance with the policies and protocols approved by the Institutional Animal Care and Use Committee (IACUC) at Washington University and all studies adhered to NIH guidelines. Mice of both sexes were used for experiments, unless otherwise indicated. Mice were kept under standard housing conditions (lights on at 6:00 am, 22 °C), with food and water available *ad libitum*. *KOR-Cre* mice^93^ harboring the coding sequence for Cre recombinase in the coding region of in exon 2 in *Oprk1*, thereby expression of functional *KOR*, were a gift from Dr. Sara Ross (University of Pittsburgh). *Vgat-flp* mice B6.Cg-*Slc32a1^tm^*^1.1^(flpo)*^Hze^*/J (JAX 029591) and *Vglut2-flp* mcie B6.129S-*Slc17a6^tm^*^1.1^(flpo)*^Hze^*/J (JAX 030212) were from the Jackson Laboratory. For HFD studies, mice were subjected to a 60% HFD (D12492, ResearchDiets) starting from 8 weeks of age. Male and female mice were included and analyzed together, unless otherwiseindicated.

### Virus and stereotaxic injections

AAV5-hSyn-DIO-hM4D(Gi)-mCherry (7.8×10^12^ vg/ml, Addgene), AAV5-hSyn-DIO-hM3D(Gq)-mCherry (7.8×10^12^ vg/ml, Addgene), AAV-retro-Ef1a-DIO-FLPo-WPRE-hGHpA (1.6×10^13^ vg/mL, Addgene), AAV1-hSyn-FLEx-mGFP-2A-Synaptophysin-mRuby (7×10¹² vg/mL, Addgene), and pGP-AAV9-syn-FLEX-GCaMP7s-WPRE 1.2×10^13^ vg/ml, Addgene) were purchased from Addgene. AAV5-EF1a-fDIO-EYFP-WPRE (4.5×10^12^ vg/ml, UNC), AAV5-Ef1a-DIO-mCherry (5.1×10^12^ vg/ml, UNC) and AAV5-hsyn-CON/Fon-EYFP (5×10¹² vg/mL, UNC) were purchased from UNC Vector Core. AAV5-EF1a-DIO-eYFP (1.8×10^13^vg/ml, Hope Center), and AAV5-CBA-DIO-GFP-Tet Tox-WPRE-bHGpA (1×10^13^ vg/ml, Hope Center) were produced by Hope Center at Washington University in St. Louis.

Briefly, mice were anesthetized in an induction chamber (4% isoflurane), placed in a stereotaxic frame (Kopf Instruments), and anesthesia was maintained with 2% isoflurane. Mice were then injected unilaterally or bilaterally according to the experimental design. Viruses were injected into the POA (AP +0.40 mm, ML ± 0.45 mm, DV -5.35 mm), DMH (AP -1.70 mm, ML ± 0.35 mm, DV -5.25 mm) using a blunt needle Neuros Syringe (65457–01, Hamilton Com.) and syringe pump (World Precision Instruments) at a rate of 30 nl/min. The needle was withdrawn 5 min after the infusion. The animal was placed on a warmed blanket until recovery from anesthesia before being returned to its home cage.

For chemogenetic activation of POA^KOR+^ neurons, KOR-Cre mice were injected bilaterally with 150 nl AAV5-hSyn-DIO-hM3D(Gq)-mCherry or AAV5-EF1a-DIO-eYFP into POA.

For chemogenetic inhibition of POA^KOR+^ neurons, KOR-Cre mice were injected bilaterally with 150 nl AAV5-hSyn-DIO-hM4D(Gi)-mCherry or AAV5-EF1a-DIO-eYFP into POA.

For chronic silencing of POA^KOR+^ neurons, KOR-Cre mice were injected bilaterally with 200 nl AAV5-CBA-DIO-GFP-Tet Tox-WPRE-bHGpA or AAV5-EF1a-DIO-eYFP into POA.

For fiber photometry recording of POA^KOR+^ neurons, KOR-Cre mice were injected unilaterally with 150 nl pGP-AAV-syn-FLEX-GCaMP7s-WPRE into POA. An optical fiber (400-μm core diameter) was then implanted 100 μm above the injection site.

For the axon terminal labeling, 200 nL AAV5-EF1a-DIO-eYFP and AAV1-hSyn-FLEx-mGFP-2A-Synaptophysin-mRuby were injected unilaterally into the POA of *KOR-Cre* mice.

For labeling GABAergic KOR neurons and non-GABAergic KOR population in the POA, mix virus of AAV5-hsyn-CON/Fon-EYFP and AAV5-Ef1a-DIO-mCherry (ratio 1:1) were injected unilaterally into the POA of *KOR-Cre*/*Vgat-flp* mice or *KOR-Cre*/*Vglut2-flp* mice.

### Anatomical tracing

For anterograde viral tracing experiments, virus AAV5-EF1a-DIO-eYFP was injected at least 6 weeks prior to transcranial perfusions with 4% paraformaldehyde to allow for anterograde transport of the fluorophore. For retrograde viral tracing experiments, after the virus AAV-retro-Ef1a-DIO-FLPo-WPRE-hGHpA was injected into DMH and AAV5-EF1a-fDIO-EYFP-WPRE into POA, there was a 3-week wait prior to perfusion to allow sufficient time for retrograde transport of the virus.

### Chemogenetic manipulations

For DREADD experiments, AAV5-hSyn-DIO-hM4D(Gi)-mCherry, AAV5-hSyn-DIO-hM3D(Gq)-mCherry or AAV5-EF1a-DIO-eYFP were injected into POA of *KOR-Cre* mice. Following a recovery period of 3 weeks, Clozapine N-oxide dihydrochloride (CNO, 1 mg/kg, Hellobio) was dissolved in saline and injected intraperitoneally to perform thermal imaging, metabolic monitor, thermal gradient assay, and GTT test. Data were discarded when viral expression was not restricted to POA or off target.

### Quantitative thermal imaging and body temperature measurements

To measure body temperatures, we used FLIR E53 thermal imaging cameras and IPTT-300 temperature transponders (Avidity Science). Quantitative thermal image videos were recorded and subsequently analyzed. In brief, one day prior to study, the interscapular region was shaved under isoflurane anesthesia. On the testing day, mice were placed in a placed in a cylindrical container (18 cm in diameter) filled with bedding and acclimated for 2 h. Following a 10 min basement recording, mice were injected with CNO and the thermal mages were collected continuously for another 2 h. Regions of interest containing BAT, eye, or a point at approximately 1 cm from the base of the tail were identified in the highest temperature at 5-min intervals. BAT temperature readings were taken at the warmest point of the intrascapular region. Eye temperature readings were taken at the warmest point of the eye. Tail temperature readings were taken ∼1 mm away from the base of the tail. For wireless temperature transponders, a DAS-8027 reader (Bio Medic Data Systems Seaford, DE) was held in close proximity under the behavioral arena and temperature from the implanted transponder was recorded.

For long term Body temperature measurements, temperature logging devices iButton (DS1925L; Analog Devices, US) were implanted in mice anesthetized with 2% isoflurane and maintained in oxygen. The iButton was inserted subdermally on the back site of the mice. The iButton was programmed to measure the body temperature every 10 min and data was collected after sacrifice. Mice were given a minimum of 7 days to recover from the procedure before starting experiments.

### Metabolic phenotyping

Metabolic phenotyping including indirect calorimetry, activity monitoring, and food consumption was carried out using a Phenomaster system (LabMaster System; TSE Systems, Bad Homburg, Germany) provided through the Washington University Nutrition Obesity Research Center. Briefly, animals were singly housed in metabolic cages in the Phenomaster with ad libitum access to food and water. Food intake was measured by subtracting the weight of uneaten food from the initial food weight. Gases were sampled from each cage for 1 min every 10 min with a flow rate of 27 L/h. Metabolic rate was measured using oxygen consumption (mL/h) and carbon dioxide production (mL/h), which was calculated via comparison of the air from the reference cage that had no animal in it. The RER was calculated by dividing CO2 production (VCO2) by O2 consumption (VO2). Energy expenditure was converted from the metabolic rate using constants CVO2 = 3.941 and CVCO2 = 1.106 and the equation EE = (CVO2 *VO2 + CVCO2 * VCO2)/1000. Physical activity was obtained by quantification of beam breaks. Activity was sampled at 100 Hz and reported in 10-min summed bins which were consolidated to 30-min bins. Z axis as well as X and Y axes movements were captured and quantified. Physical activity is reported as total beam breaks for all three axes by the animals in the cage. For chemogenetic activation and inhibition manipulation, animals were habituated for 3 days to Phenomaster housing. On day 4, at 10:00 a.m., animals were injected peritoneally with Clozapine N-oxide dihydrochloride (CNO, 1 mg/kg, Hellobio) and the metabolic rate was acquired. Then the mice were allowed 4∼5 days recovery before the next CNO injection at 6 pm timepoint. For chronic silencing experiments, mice were put in the metabolic chamber 1 day after the surgery and monitored continuously for about 2 weeks.

### Thermal gradient test

Mice were placed in a thermal gradient test apparatus (GNF Systems Gradient Plate v.1.0; GNF Systems, San Diego, CA) ranging in temperature from ∼12 to ∼48°C as described previously^94^. The thermal chamber (100 cm × 7.6 cm × 10cm) creates a thermal gradient spited into 20 thermal zones and mice were allowed to freely explore the chamber to select the optimal-temperature position. Mice cages were placed in the test room for acclimation overnight prior to the formal test. Experiments were recorded using a video camera, and the location of the mice was automatically detected using Bonsai software. Before the test, mice were injected with CNO (i.p., 1 mg/kg) diluted in sterile saline and data from a total of 2h recording session were used for analysis. Temperature preference was defined as the temperature of the zone in which a mouse spent the most time for the entire test.

### Body composition analysis

For body composition measurement, fat and lean mass of each mouse in this study were measured by an EchoMRI-100H, quantitative nuclear resonance system (EchoMRI LLC). For chronic silencing experiments, the body composition analysis was measured one week before surgery and 2 or 3 weeks after virus injection.

### Quantification of feeding and activity in the home cage

We used FED3 devices to monitor feeding of singly housed mice. The FED3 devices were programmed to dispense one 20-mg standard chow pellet (Testdiet) at a time and automatically replaced the pellet after it was taken. Timestamps for each dispensed pellet were recorded. FED3 offered greater precision than the metabolic phenotyping home cages^95^. We use free feeding code program to dispense 20 mg chow pellets and record the time each pellet is taken in “free feed mode”. The FR1 code program was used to dispense pellets when mice interact with FED3 through two nose-pokes and FED3 responds to the mice by delivering pellets whenever mice poke the active port. The FED3 was the only source of food in the cage. Mice were given one week to habituate to FED3 feeding prior to experiments. Stable feeding was required before experiments. Mice and functioning of the FEDs were checked daily. Pallidus, a wireless home cage sensor was used to monitor locomotor activity, light, and ambient temperatures in the home cage. Both the FED3 and Pallidus device were designed and developed by Dr. Alexxai Kravitz at Washington University School of Medicine.

### Glucose-tolerance tests

GTTs were performed after a fasting period of 6 hours. Blood glucose concentrations were measured from tail blood using an automatic glucose monitor (Bayer HealthCare Ascensia Contour). Each mouse received an intraperitoneal injection of glucose (50% Dextrose, Hospira, IL) at a dose of 2.0 g/kg body mass in HFD study. Blood glucose concentrations were measured at baseline and after 15, 30, 60, 90 and 120 min.

### Warm temperature exposure

For thermal challenges, the mice in the warm condition were placed in a clean cage wrapped by a circulating water blanket which was set to 38°C. Mice in the room temperature condition were placed in a clean cage in a 22–23°C room. Water was supplied ad libitum in all cages. Cages in the warm condition were given enough time to reach the target temperature as confirmed by a thermometer before mice were placed inside of them. Temperature exposures lasted for 4 hr, after which mice were immediately anesthetized with pentobarbital and transcardially perfused with 4% paraformaldehyde in phosphate buffer, and brains were subsequently collected.

### Immunohistochemistry

IHC was performed as previously described by^62^. In brief, mice were intracardially perfused with 4% PFA and then brains were sectioned (30 microns) and placed in 1×PB until immunostaining. Free-floating sections were washed in 1× PBS for 3 × 10 min intervals. Sections were then placed in blocking buffer (0.5% Triton X-100% and 5% natural goat serum in 1× PBS) for 1 hr at room temperature. After blocking buffer, sections were placed in primary antibody rabbit Phospho-c-Fos (Ser32) antibody (RRID:AB_10557109, 1:500 Cell Signaling Technology) overnight at room temperature. After 3 × 10 min 1×PBS washes, sections were incubated in secondary antibody goat anti-rabbit Alexa Fluor 633 (RRID:AB_2535731, 1:1000, Invitrogen) for 2 hr at room temperature, followed by subsequent washes (3 × 10 min in 1× PBS then 3 ×10 min 1× PB washes). After immunostaining, sections were mounted on Super Frost Plus slides (Fisher) and covered with Fluoromount-G mounting medium with DAPI (Invitrogen) and cover glass prior to being imaged on a Leica DM6 B microscope. Images were analyzed by an observer blinded to the group or genotype using Image J software from NIH Image (version 1.34e).

### Brain clearing and light sheet microscopy

Whole brain clearing was performed using electrophoretic transport-based CLARITY^96^, using SHIELD tissue preservation^97^. Briefly, mice were transcardially perfused with ice-cold PBS, followed by 4% paraformaldehyde. Samples were post-fixed in 4% PFA overnight and were rinsed using PBS. Brains were infused with SHIELD-off polyepoxy (LifeCanvas Biotechnology, Cambridge, MA) for 4 days and polymerized in SHIELD-ON solution at 37°C for 24 hours. Putative delipidation was performed at 45°C for 24 hours, and samples were actively cleared using a SmartBatch+ ETC platform for 30 hours (LifeCanvas). Cleared brains were washed in PBS for 24 hours, and were refractive index matched to 1.52 using EasyIndex refractive index matching solution (RIMS, LifeCanvas Biotechnology). Whole brain imaging was performed on a Miltenyi Ultra Microscope Blaze lightsheet platform using a 4x/0.35 objective. Images were acquired on ImSpector 7.7 (Miltenyi biotech), stitched with Stitchy 1.15 (Translucence Biotechnology), and visualized in Imaris 10.2 (Oxford Instruments).

### Fluorescent in situ hybridization (FISH)

Following rapid decapitation of mice, brains were flash frozen in -50°C 2-methylbutane and stored at -80°C for further processing. Coronal sections containing the POA region, corresponding to the injection plane used in the behavioral experiments, were cut at 20 mM at - 20°C and thaw-mounted onto Super Frost Plus slides (Fisher). Slides were stored at -80°C until further processing. FISH was performed according to the RNAScope 2.0 Fluorescent Multiple Kit User Manual for Fresh Frozen Tissue (Advanced Cell Diagnostics, Inc) as described^98^. Slides containing the specified coronal brain sections were fixed in 4% paraformaldehyde, dehydrated, and pretreated with protease IV solution for 30 min. Sections were then incubated for target probes for mouse *Oprk1* (Oprk1, accession number NM_001204371.1, probe region 256-457) and *Vgat* (Slca32a1, NM_009508.2, probe region 894-2037) for 2 hr. All target probes consisted of 20 ZZ oligonucleotides and were obtained from Advanced Cell Diagnostics. Following probe hybridization, sections underwent a series of probe signal amplification steps (AMP1–4) including a final incubation of fluorescently labeled probes (Alexa 488, Atto 550, Atto 647), designed to target the specified channel (C1–C3 depending on assay) associated with the probes. Slides were counterstained with DAPI and coverslips were mounted with Fluoromount-G mounting medium with DAPI (Invitrogen). Alternatively, mice transcardially perfused with cold PBS and PFA with fixed brain tissue collected and sectioned at 30 mM as described previously were processed for FISH as above. Images were obtained on a Leica DM6 B upright microscope (Leica), and Application Suite Advanced Fluorescence (LAS AF) and ImageJ software were used for analyses. To analyze images for quantification of *Oprk1*/*Vgat* coexpression, each image was opened in ImageJ software, channels were separated, and an exclusive fluorescence threshold was set. We counted total pixels of the fluorescent signal within the radius of DAPI nuclear staining, assuming that each pixel represents a single molecule of RNA as per manufacturer guidelines (RNAscope). A positive cell consisted of an area within the radius of a DAPI nuclear staining that measured at least five total positive pixels. Positive staining for each channel was counted in a blind-to-condition fashion using ImageJ or natively in LAX software (Leica).

### Adipose tissue histology

Mice were deeply anesthetized with isoflurane, and BAT, inguinal white adipose tissue (iWAT), and epididymal white adipose tissue (eWAT) were dissected, fixed in 4% PFA overnight, embedded in paraffin, and sectioned at 5 μm thickness. Histological changes were examined after hematoxylin and eosin (H&E) staining and brightfield images were obtained with Hamamatsu NanoZoomer HT 2.0 slide scanning system. The measurement of adipose cell size was performed using ImageJ.

### Western blot

Western blotting was performed as previously described^99^. Briefly, ∼50-80 mg of tissue was homogenized in RIPA lysis buffer (Cell Signaling Technology) with added protease and phosphatase inhibitors (Cell Signaling Technology), using a PolytronTM handheld homogenizer on ice. Samples were centrifuged at 12000 x g, for 10 minutes, at 4°C. The supernatant was removed to a new tube, being careful to avoid the floating lipid layer. Lysates were normalized to protein concentration, determined using the DC protein assay (Bio-Rad), and combined with NuPAGETM LDS Sample Buffer (ThermoFisher), then heated at 37°C for 10 minutes. Samples were run on 4-12% Bolt™ Bis-Tris protein gels (ThermoFisher) with MOPS buffer (ThermoFisher) then transferred to Immobilon-FL PVDF (Sigma). Membranes were placed in blocking buffer (5% nonfat dried milk in TBS) for 1 hour at room temperature, then placed in primary antibodies overnight at 4°C, and images were obtained and quantified using the Licor system with Image Studio Lite software. All primary antibodies were diluted in TBST (TBS with 0.1% Tween®20) containing 5% bovine serum albumin. Antibodies and dilutions used were: OXPHOS Rodent cocktail (Abcam #ab110413, 1:500), Phospho-HSL Ser660 (CST #45804, 1:1000), HSL (CST #18381, 1:1000), Phospho-PLIN Ser522 (Vala-Sciences #4856, 1:5000), PLIN (CST #9349, 1:1000), and Vinculin (CST #13901, 1:1000). Western blots were quantified by Image J.

### Fiber photometry recording

For all fiber photometry experiments, the same strategy to selectively label POA^KOR+^ neurons was used. *KOR-Cre* mice were injected unilaterally with AAV-syn-FLEX-GCaMP7s-WPRE in POA. Simultaneously, mice were implanted with fiber-optic cannulas (200 μm) in POA (D/V -5.20). Mice that received GCaMP7s injection into the POA were allowed 3 weeks for recovery, then these mice were acclimatized to a custom-made plastic rectangular testing chamber (28 cm x 31 cm x 32 cm) filled with bedding for 24 hours before experiments. Recording of Ca^2+^ dependent and isosbestic signals were obtained using previously described methods with Bonsai software and FP3002 (Neurophotometrics)^100^. 470 and 415 nm LEDs were used to record interleaved isosbestic and Ca^2+^ dependent signals following the manufacturer directions. Fiber photometry adopted a sampling rate of 4 Hz^65^. Videos were recorded at 4 fps.

For long-term continuous recording of POA^KOR+^ neurons under normal feeding, fasting and refeeding, time constrained feeding, light shifts or wheel running activity, mice were single housed in the testing chamber with free access to automated FED3 device and cotton bedding and allowed 24 hours acclimatization to the cage. We then tethered the freely moving mice through their ferrule implants to the fiberoptic cables (200-μm core, NA = 0.37, bundled fibers, Neurophotometrics, Inc, now MBF Bioscience) using black ceramic mating sleeves (Thorlabs). We use a 10% LED duty cycle, low power output, with a 25ms period and sample at 4 Hz sampling rate with using FP3002 (Neurophotometrics) and Bonsai software including light intensity for 470 wavelength was 15 uW and 25 uW of total light at the tip of the fiber patch cord. Normal feeding was monitored using FED3 devices dispensing 20 mg chow pellets at each time for continuous 24 hours. Fasting and refeeding was recorded by removing food access for 18 hours during recording of fiber photometry signals. Time constrained feeding only restricts feeding to one out of every three hours to consolidate feeding times without limiting food intake. Light shifts were achieved by applying 6 light hours in advance. A rodent running wheel was put in the testing chamber to record wheel running activities.

For temperature challenges, the animal was placed in a 32 x 57 x 45.5 cm temperature plate (BIO-T2CT, Bioseb) and enclosed by Plexiglas chamber walls. Animals were allowed to recover after patch cord attachment for at least 30 min. For warm temperature stimuli, the floor temperature was changed from 22°C to 34°C, while 22°C to 10°C was used for cool stimuli.

For fiber photometry data analysis, a custom Python script was used to fit the isosbestic signal (415 nm) to the calcium-dependent signal (470 nm) and correction for photobleaching using biexponential curve fitting. Deinterleaved signals were analyzed using methods as previously reported^101^. Briefly, raw signals were smoothed using a moving average, fitted with an exponential curve using a non-linear least squares function for baseline correction, signals were standardized using the mean value and standard deviation (Z-Score), the standardized isosbestic and Ca^2+^ signals were scaled a non-negative robust linear regression, and normalized ΔF/F was calculated. The Ca^2+^ events were defined by a rise of two standard deviations above baseline. Fasted mice (24 h) that showed >10% ΔF/F signal decrease after food presentation were selected for further analysis (∼60% of total injections). We also plotted these Ca^2+^ events by using Rayleigh plots to examine time-of-day dependent changes. Events on the Rayleigh plots are Ca^2+^ events which were defined by a rise of two standard deviations above baseline. All time points were calculated as a radian and plotted for that specific time event during a 24 hour time. We used findPeaks, a python library for signal processing to find peaks in the continuous photometry Z-Score. A peak was defined as a point in the data where the signal is higher than the neighboring points. Additionally, we included the parameter prominence of 2.5, which measures the vertical distance between the peak and its lowest contour around its surrounding baseline.

For the recorded specific behaviors, Bonsai scripts were used to track animals, to time stamp changes in regions of interests, such as their nest area, and to quantify velocity of movement. We also analyzed the Ca^2+^ transients and Ca^2+^-dependent signals in relation to periods of movement, feeding, wheel running, and time in the nest (or other areas). Food intake rhythms were collected using automated FED3 feeding device and were converted to 1-hour bins and analyzed with a custom Python script.

Nestlet entrance and exit was quantified by using Bonsai by drawing a region around the animal’s nestlet location in the arena after habituation. Live acquisition using background subtraction and centroid tracking recorded when the animal centroid arrives or leaves the nestlet region, the amount of time including the number of entries and exits were timestamped. For peak analysis timestamped peak events were aligned to the timestamped events of the animal being inside/outside of the nestlet region.

Pellet retrieval was timestamped via TTL pulse generated from an Arduino microcontroller board that was communicating with Bonsai via StandardFirmata. A feeding bout was defined by calculating the time difference between pellet retrievals. It was established that 30 seconds was the time threshold for a feeding bout including at least 3 pellets retrieved by the animal considering a 3 second timeout after each pellet retrieval. A custom python script was used to calculate and align to fiber photometry Z-Score for peri-event analysis.

For wheel running experiments, a custom made 5 inch mouse exercise wheel was attached to a 3D-printed mouse running wheel stand. A magnet was glued to the mouse wheel and a hall sensor placed inside a cavity of the stand. Each revolution was defined when the magnet disrupted the hall sensor magnetic field then a timestamped TTL pulse was generated from Arduino that was communicating with Bonsai via StandardFirmata. Wheel running bouts were defined by calculating the time difference between wheel revolutions. It was established that the threshold was 4 seconds and at least 15 rotations. A custom python script was used to calculate and align to fiber photometry Z-Score.

At the end of recordings, mice were processed for post hoc histological evaluation.

Animals were excluded when the virus or the fiber tip was off target.

For ΔF/F calculation, baseline F0 was defined as the average F during 4 min before feeding behavior was initiated in the fasting induced calcium activity recording experiment, and F0 was defined as the average F during 5 min before the ambient temperature was changed in the temperature induced calcium activity recording experiment.

### Statistical analysis

Statistical analysis was carried out using Prism software (GraphPad; San Diego, CA). For comparisons between two sets of time-course data, different Student’s t tests were used. For experiments where the same mice were tested in multiple conditions a paired test was applied, and when different mice were compared an unpaired test was used. For time-course thermal data p values were not corrected for multiple comparisons across time. For comparisons of three or more groups, a one-way ANOVA was used. Repeated measures ANOVA tests were used for experiments where the same subject was used across multiple conditions. For each experiment, control groups and statistics are described in the main text. For fiber photometry statistical analysis, the mean signal of the baseline and stimulus windows was used, and comparisons were made using the Wilcoxon ranked-sum test, with α=0.95. All ‘n’ values represent the number of animals in a particular group for an experiment. All data are reported as mean ± SEM.

---

**Extended Data Fig. 1.**
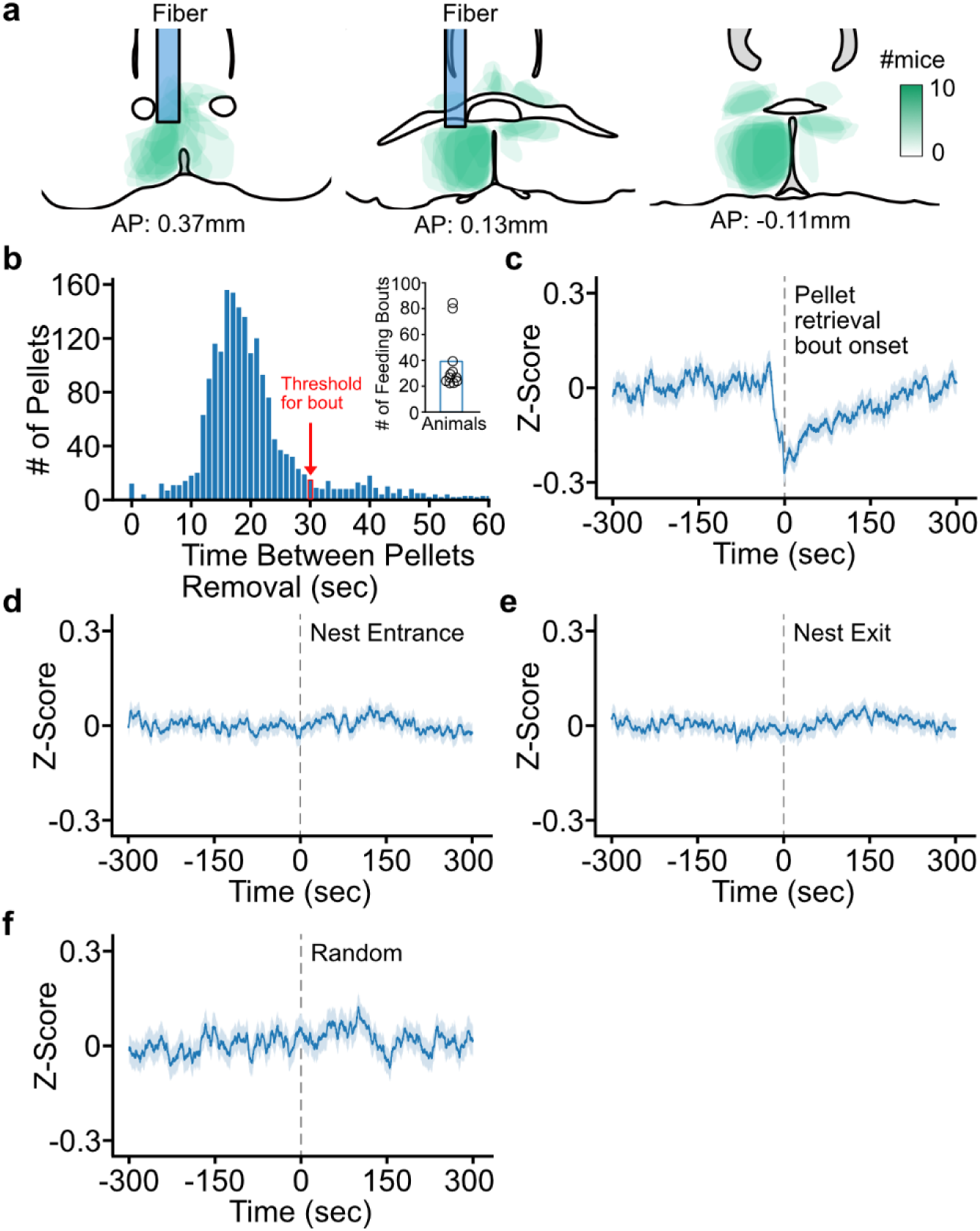
Characterization of POA^KOR+^ neurons fiber photometry recordings under *ad libitum* feeding. **a**, Heatmaps of Gcamp7s-GFP expression and fiber placement at different Bregma sites from all experimental mice. The relative scale for the expression intensity (region infected by fluorescence coverage) was shown in the right. **b**, Distribution of times between pellet removals during the 24-hour recording period. Right: Feeding bouts cut at within 30 seconds of the prior pellet during the 24-hour window. **c**, Ca^2+^ signals of the onset of a feeding bout during *ad libitum*. **d**, POA^KOR+^ neuron fiber-photometry recording of mice entering the nest area. **e**, POA^KOR+^ neuron fiber-photometry recording of mice exiting the nest area. **f**, Ca^2+^ signals of the random movement. *P < 0.05, **P < 0.01, ***P < 0.001, and ****P < 0.0001; ns, no significance. n=12 mice per group. Data are means ± s.e.m.

**Extended Data Fig. 2.**
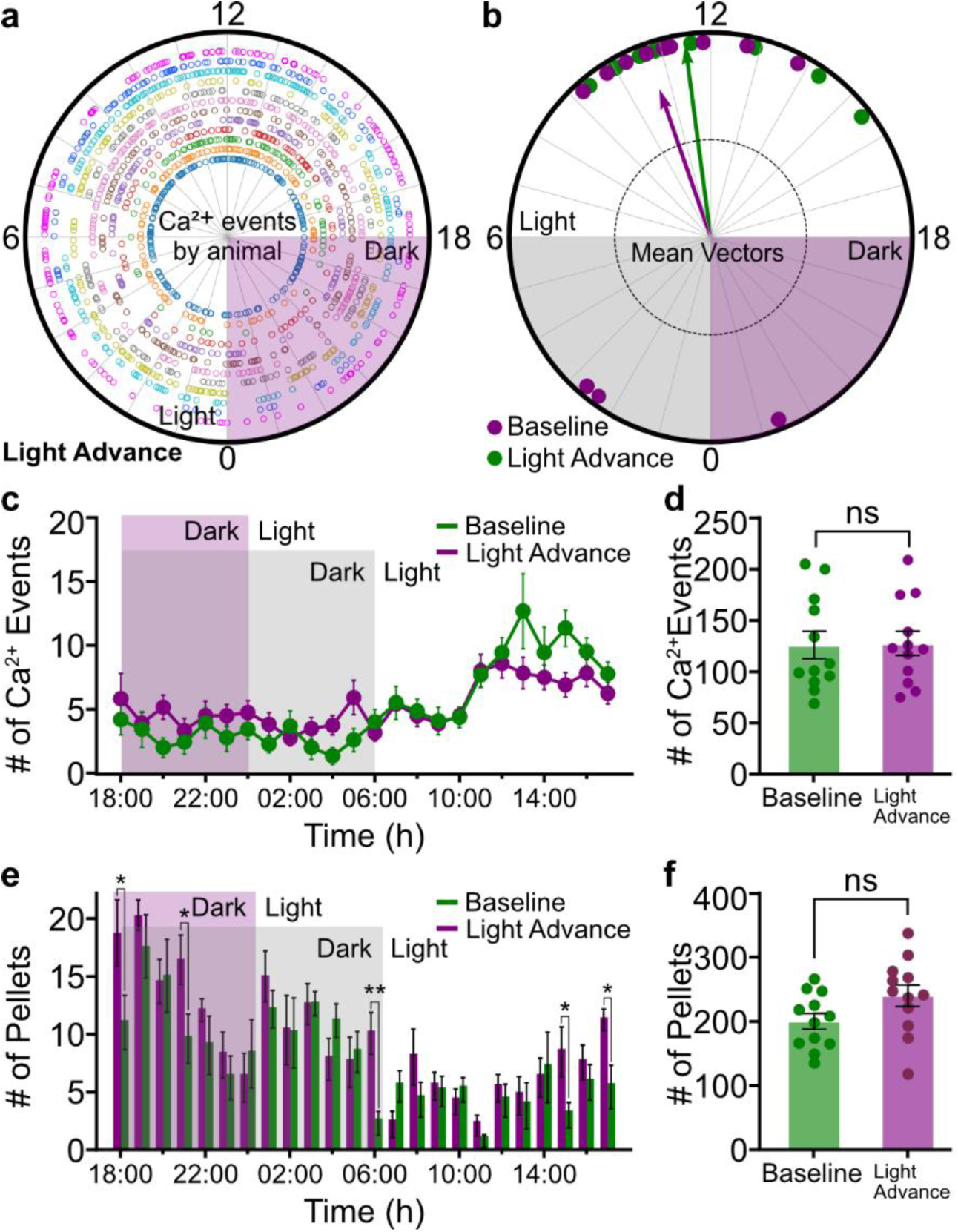
POA^KOR+^ neurons activity is not altered to a phase shift. **a**, Raleigh plots of calcium events by individual animal under the light advance shift scheme. **b**, Mean Raleigh vector of calcium events distribution under the light advance shift scheme. **c**, Distribution of calcium events per hour of POA^KOR+^ neurons under the light advance shift. **d**, Comparison of number of calcium events per hour recorded between normal light-dark circle (Baseline) and light advance shift condition. **e**, Comparison of number of food pellets taken per hour recorded under normal light-dark circle and light advance shift condition. **f**, Comparison of number of food pellets taken under normal light-dark circle (Baseline) and light advance shift condition. Rayleigh test for non-uniformity of circular data and circular one-way ANOVA to compare circular data, two-way ANOVA comparing baseline and light advance events and pellets in (**c**) and (**e**), unpaired-sample *t* test in (**d**) and (**f**). *P < 0.05, **P < 0.01, ***P < 0.001, and ****P < 0.0001; ns, no significance. n=12 mice. Data are means ± s.e.m.

**Extended Data Fig. 3.**
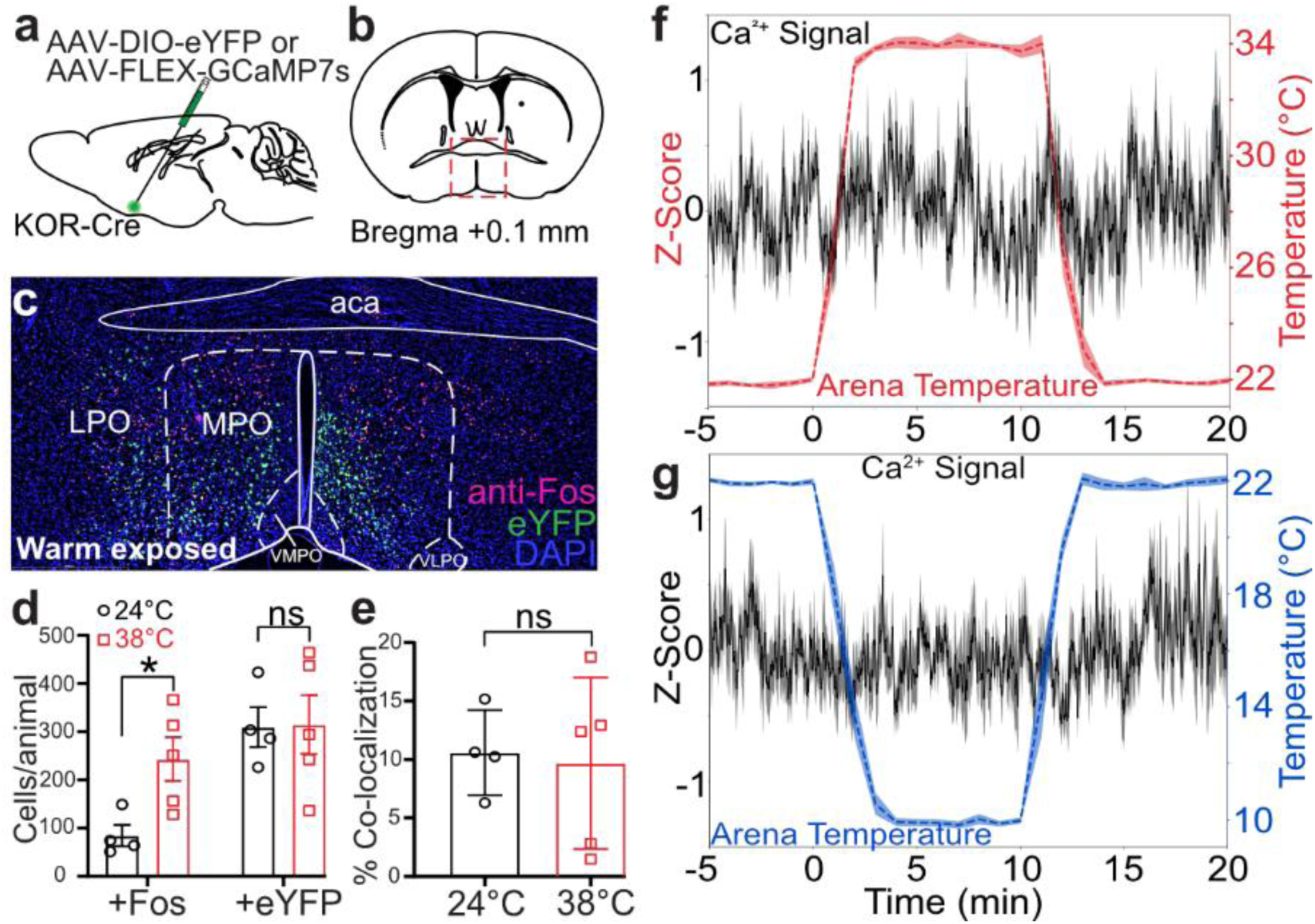
POA^KOR+^ neurons are not temperature-sensitive neurons. **a**, Schematic of virus injections of AAV-FLEX-GCaMP7s or AAV-EF1a-DIO-eYFP into the POA of KOR-Cre mice. **b**, Representative Bregma section showing targeted POA region. **c**, Representative images of Fos expressing neurons (red) and KOR neurons (green) from animals exposed to warm temperature stimuli at 38°C. **d**, Quantification of Fos positive neurons under warm (38°C) or room temperature (24°C) in KOR positive neurons or non-KOR positive neurons. **e**, Quantification of overlapping of both Fos positive neurons and KOR positive neurons under warm (38°C) or room temperature (24°C). **f**, **g**, Fiber photometry recordings of POA^KOR+^ neurons respond to warm (**f**, 34°C) or cool temperature (**g**, 10°C). Scale bars, 250 µm. Unpaired-sample *t* test in (**d**) and (**e**), *P < 0.05, **P < 0.01, ***P < 0.001, and ****P < 0.0001; ns, no significance. n=4-5 mice per group. Data are means ± s.e.m.

**Extended Data Fig. 4.**
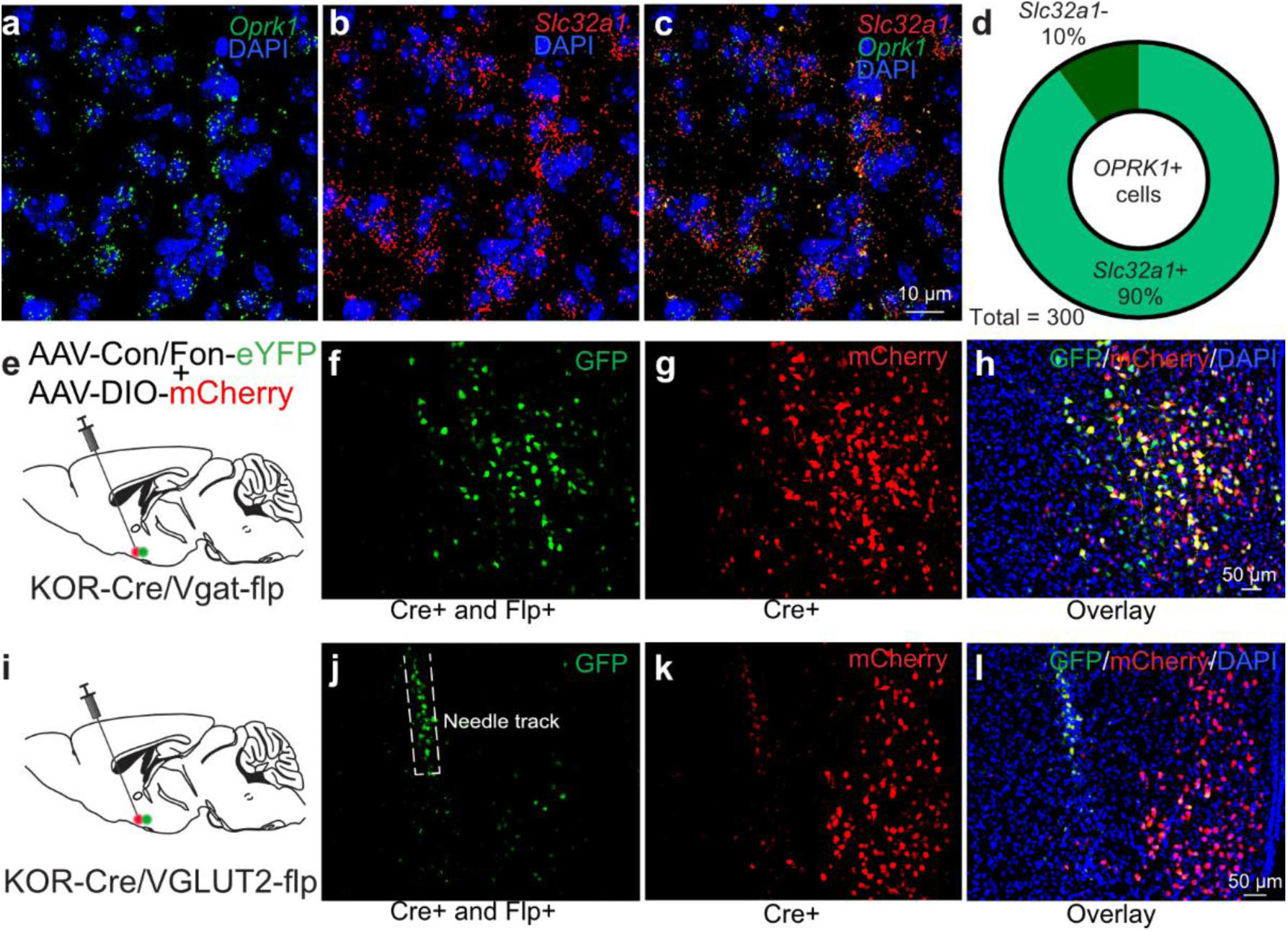
POA^KOR+^ neurons are mainly inhibitory neurons. **a**-**c**, Representative FISH images of (**a**) *Oprk1* gene which encodes KOR, *Slc32a1* (**b**), and overlays (**c**) in the POA. **d**, Quantification of cells labeled with *Oprk1* probe and *Slc32a1* probe. Scale bars, 10 µm. n=3 mice. **e**, Schematic of virus injections of AAV-hsyn-CON/Fon-EYFP and AAV-DIO-mCherry (ratio 1:1) into the POA of KOR-Cre/Vgat-flp mice to label population that are GABAergic KOR neurons. **f-h**, Representative images showing GABAergic KOR positive neurons (**f**, green), KOR positive neurons (**g**, red) and overlays (**h**) in the POA. **i**, Schematic of virus injections of AAV-hsyn-CON/Fon-EYFP and AAV-DIO-mCherry (ratio 1:1) into the POA of KOR-Cre/Vglut2-flp mice to label population that are Glutamatergic KOR neurons. **j-l**, Representative images showing Glutamatergic KOR positive neurons (**j**, green), KOR positive neurons (**k**, red) and overlays (**l**) in the POA. Scale bars, 50 µm. n=2 mice.

**Extended Data Fig. 5.**
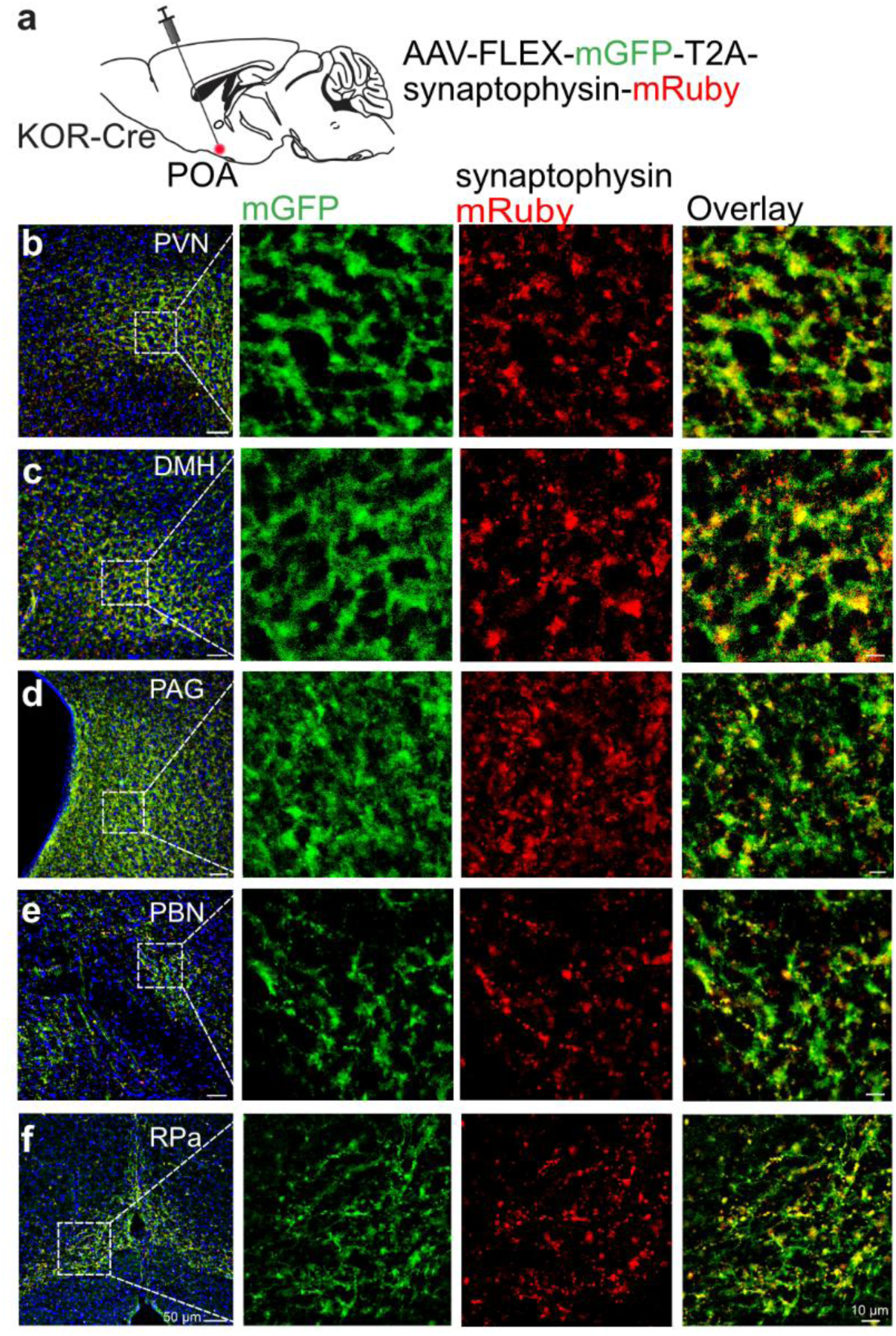
Characterization of POA^KOR+^ neurons synaptic terminals related to downstream metabolism regulation circuits. **a**, Schematic of viral injections of AAV-hSyn-FLEx-mGFP-2A-Synaptophysin-mRuby into the POA of KOR-Cre mice to label presynaptic terminals of POA^KOR+^ neurons. **b**-**f**, Major efferent projection targets POA^KOR+^ neurons as visualized with mRuby-positive puncta (red) and mGFP positive axons (green) in the (**b**) PVH, (**c**) DMH, (**d**) PAG, (**e**) PBN, and (**f**) RPa. Left panel indicates the low power images. Right panels indicate the magnified images. DMH, dorsomedial hypothalamus; DR, dorsal raphe; LC, locus coeruleus; PBN, parabrachial nucleus; PAG, periaqueductal grey; PVH, paraventricular hypothalamus; POA, preoptic area; RPa, raphe pallidus. Scale bars, 50 µm (left) or 10 µm (right). n=2 mice.

**Extended Data Fig. 6.**
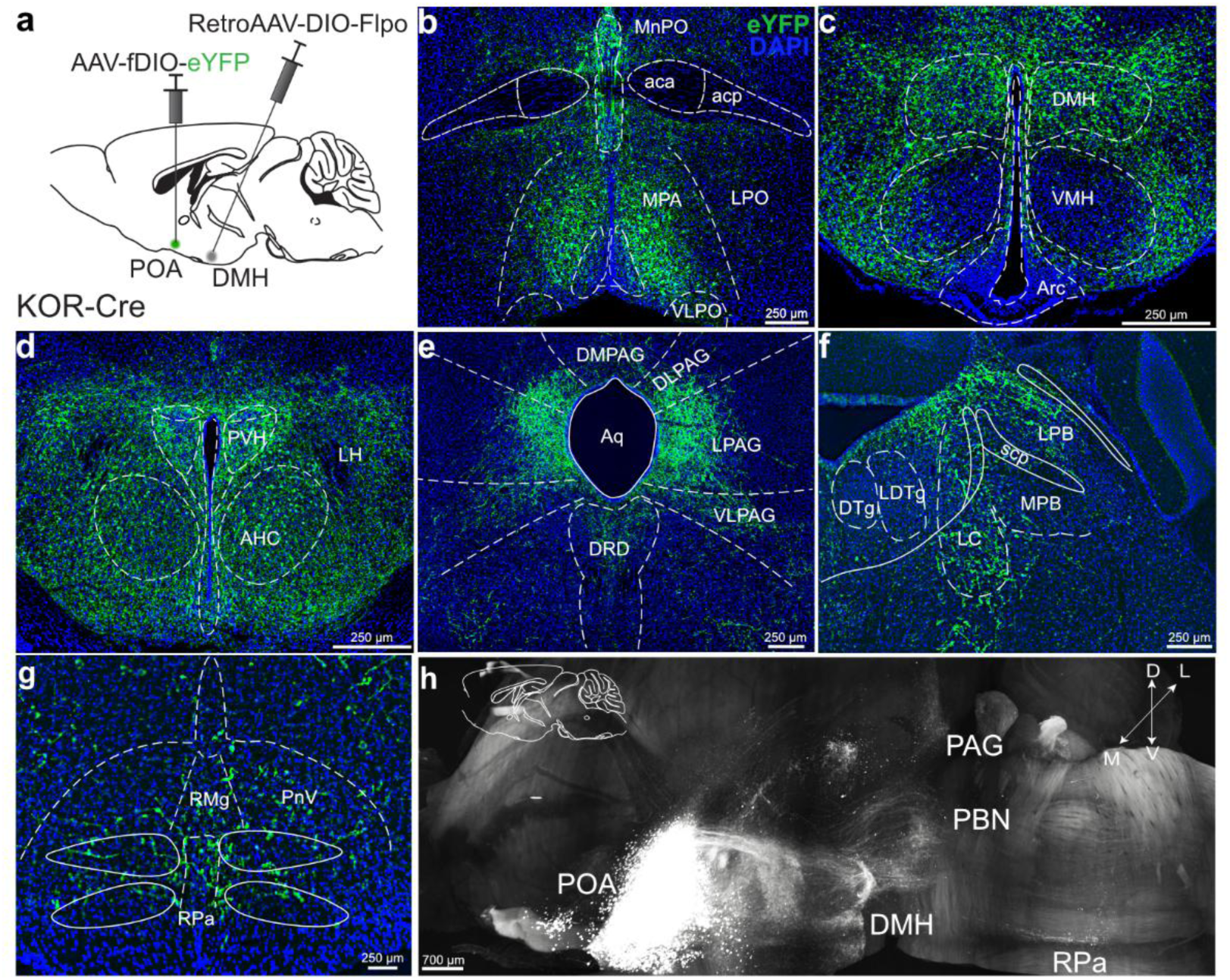
Arborization from DMH projecting POA^KOR+^ neurons share similar projection patterns compared to POA^KOR+^ neurons population. **a**, Schematic of viral injections of Retro-AAV-DIO-Flpo into the DMH and AAV-fDIO-eYFP into the POA of KOR-Cre mice to label only POA^KOR+^ neurons that project to the DMH. **b**, Cell bodies labeled with eYFP are limited to the medial side of POA. **c**-**g**, Distribution of arborizing processes from the DMH projecting POA^KOR+^ neurons are seen in multiple brain regions that share the projection patterns as local POA^KOR+^ neurons. GFP-positive fibers are seen in multiple brain regions that are implicated in the regulation of body temperature and energy expenditure including the (**c**) DMH, (**d**) PVH, (**e**) PAG and DR, (**f**) PBN and LC, and (**g**) RPa. (**h**) Summary of projection regions of POA^KOR+^ neurons. Images obtained from light sheet microscopy scanning. Major arborization regions are indicated. DMH, dorsomedial hypothalamus; DR, dorsal raphe; LC, locus coeruleus; PBN, parabrachial nucleus; PAG, periaqueductal grey; PVH, paraventricular hypothalamus; POA, preoptic area; RPa, raphe pallidus. Scale bars, 250 µm. n=3 mice.

**Extended Data Fig. 7.**
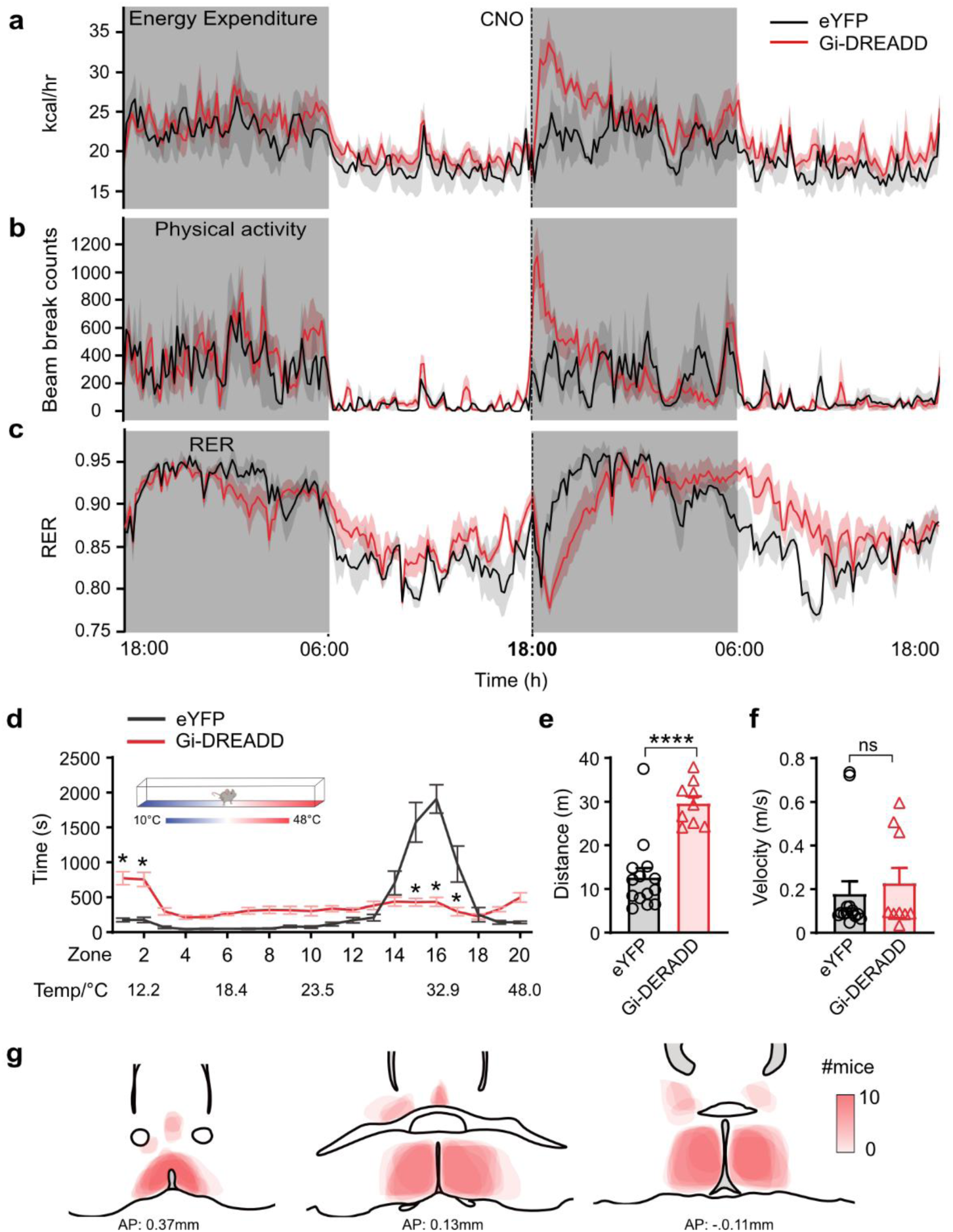
Chemogenetic inhibition of POA^KOR+^ neurons regulates energy expenditure at dark phase. **a**, **b**, Chemogenetic inhibition of POA^KOR+^ neurons after CNO (1 mg/kg, i.p.) injection at 6:00 pm right before entering the dark phase also increases EE (**a**) and locomotor activity (**b**). **c**, Chemogenetic inhibition of neurons suppresses RER. **d**, Thermal gradient assay showing the time of KOR-Cre mice staying on a different temperature zone following CNO injection. **e**, **f**, Total distance (**e**) and maximum velocity (**f**) in the thermal gradient device during the recording period. **g**, Heatmaps of Gi-DREADD expression at different Bregma sites from all experimental mice. The relative scale for the expression intensity (region infected by fluorescence coverage) was shown in the right. Statistical analyses included Two-way ANOVA (**a-d**) comparing control and treated groups after CNO injection (n=3-5 mice per group). Unpaired-sample *t* test in (**e**) and (**f**), *P < 0.05, **P < 0.01, ***P < 0.001, and ****P < 0.0001; ns, no significance. n=9-14 mice per group. Data are means ± s.e.m.

**Extended Data Fig. 8.**
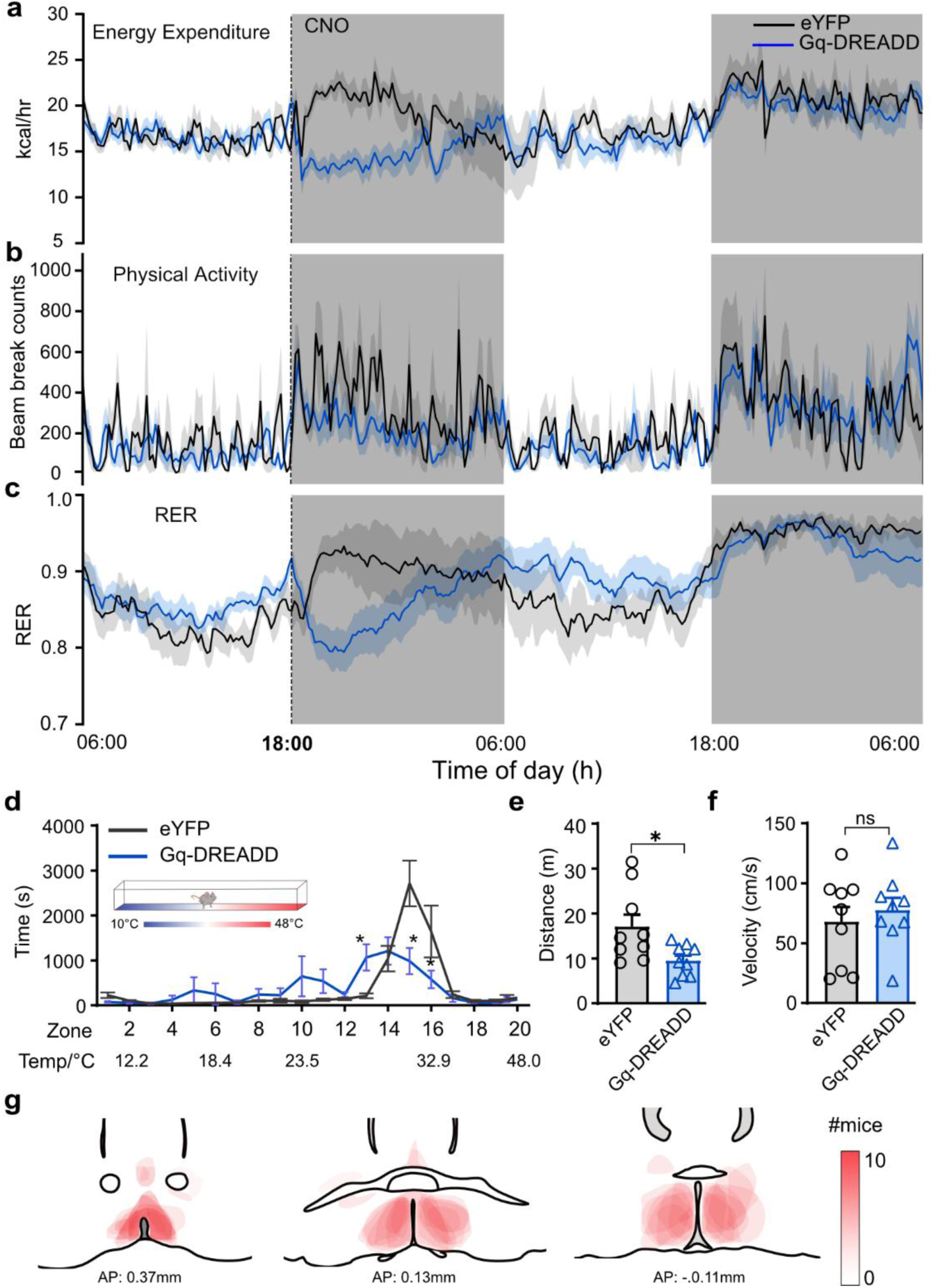
Chemogenetic activation of POA^KOR+^ neurons suppresses energy expenditure at dark phase. **a**-**c**, Chemogenetic activation of POA^KOR+^ neurons after CNO (0.5 mg/kg, i.p.) injection at 6:00 pm right before entering the dark phase also suppresses EE (**a**), locomotor activity (**b**), and RER (**c**). **d**, Thermal gradient assay showing the time of KOR-Cre mice staying on a different temperature zone following CNO injection. **e**, **f**, Total distance (**e**) and maximum velocity (**f**) in the thermal gradient device during the recording period. **d**, Heatmaps of Gq-DREADD expression at different Bregma sites from all experimental mice. The relative scale for the expression intensity (region infected by fluorescence coverage) was shown in the right. Statistical analyses included Two-way ANOVA (**a-d**) comparing control and treated groups after CNO injection. Unpaired-sample *t* test in (**e**) and (**f**), *P < 0.05, **P < 0.01, ***P < 0.001, and ****P < 0.0001; ns, no significance. n=7-9 mice per group. Data are means ± s.e.m.

**Extended Data Fig. 9.**
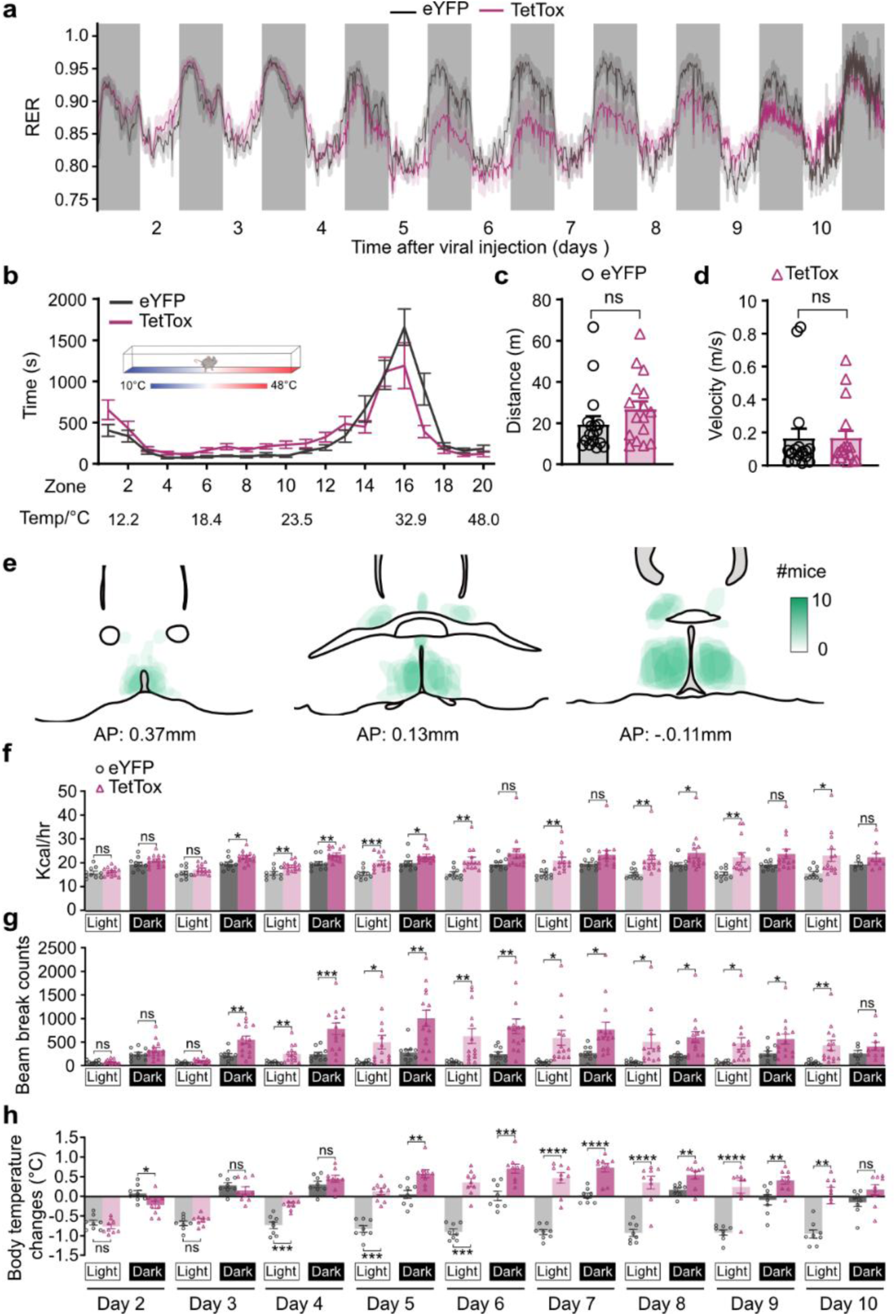
Synaptic inactivation of hypothalamic KOR neurons regulates energy expenditure and body temperature. **a**, RER decreased after silencing POA^KOR+^ neurons. **b**, Thermal gradient assay showing the time of KOR-Cre mice staying on a different temperature zone after silencing KOR neurons. **c**, **d**, Total distance (**c**) and maximum velocity (**d**) in the thermal gradient device during the recording period. **e**, Heatmaps of TetTox-GFP expression at different Bregma sites from all experimental mice. The relative scale for the expression intensity (region infected by fluorescence coverage) was shown in the right. **f**-**h**, Bar plots of EE (**f**), physical activity (**g**) and body temperature (**h**) in light and dark phase of lean KOR-Cre mice after TetTox injection. Statistical analyses included Two-way ANOVA comparing control and treated groups in (**a**) and (**b**). Unpaired-sample *t* test in (**c**) and (**d**). *P < 0.05, **P < 0.01, ***P < 0.001, and ****P < 0.0001; ns, no significance. n=10-16 mice per group. Data are means ± s.e.m.

**Extended Data Fig. 10.**
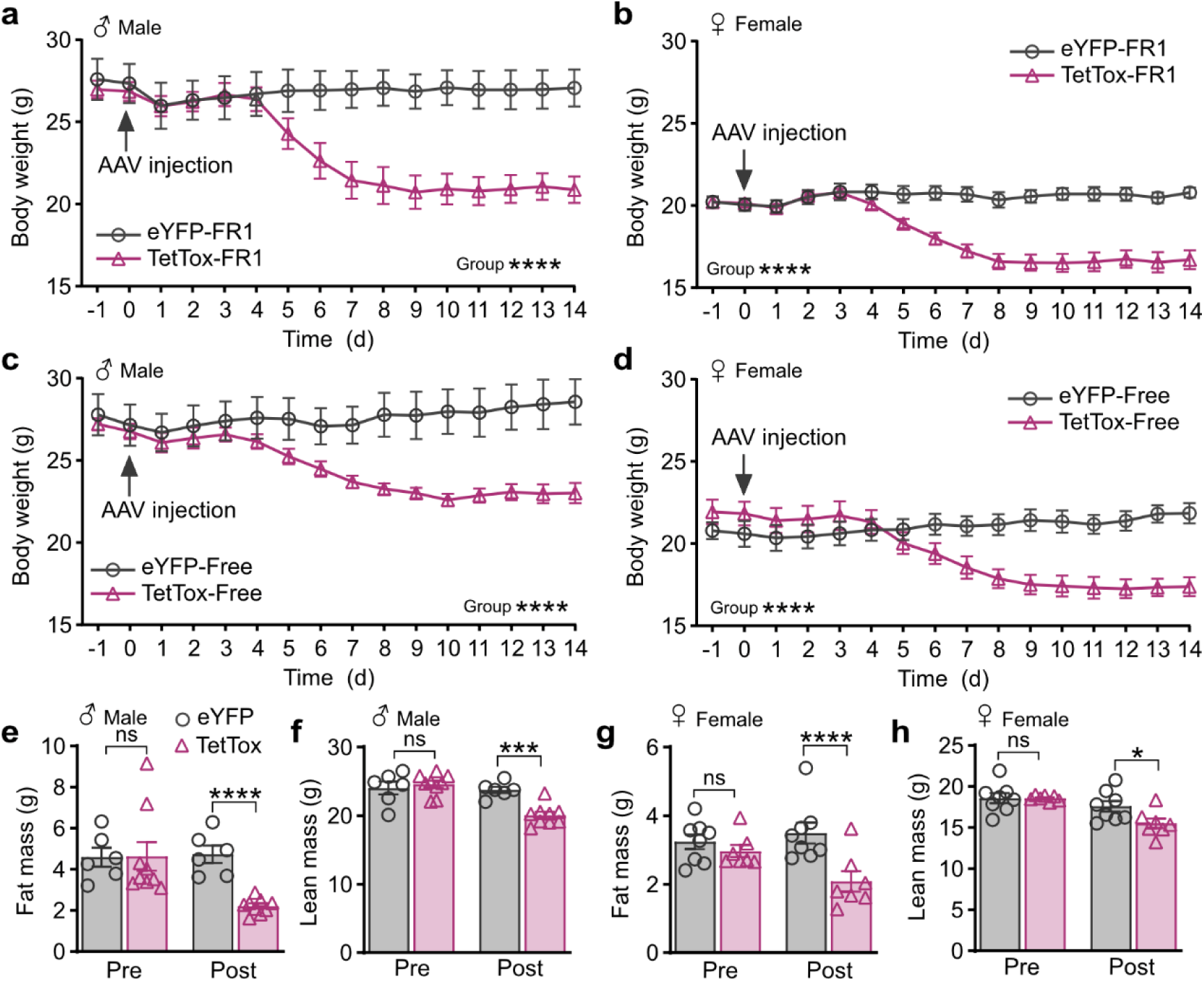
Sex variance on body weight and body mass after synaptic inactivation of hypothalamic KOR neurons. **a**, Absolute body weight changes of male KOR-Cre mice after silencing POA^KOR+^ neurons monitored with FR1 feed code. **b**, Absolute body weight changes of female KOR-Cre mice after silencing POA^KOR+^ neurons monitored with FR1 feed code. **c**, **d**, Absolute body weight changes of male (**c**) or female (**d**) KOR-Cre mice after silencing POA^KOR+^ neurons monitored with free feed code. **e**, **f**, Fat (**e**) and lean (**f**) masses of male KOR-Cre mice after silencing POA^KOR+^ neurons were measured by EchoMRI at two weeks after viral injection. **g**, **h**, Fat (**g**) and lean (**h**) masses of female KOR-Cre mice after silencing POA^KOR+^ neurons were measured by EchoMRI at two weeks after viral injection. Statistical analyses included Two-way ANOVA comparing control and treated groups in (**a**), (**b**), (**c**), and (**d**). Unpaired-sample *t* test in (**e**), (**f**), (**g**), and (**h**). *P < 0.05, **P < 0.01, ***P < 0.001, and ****P < 0.0001; ns, no significance. n=6-9 mice per group. Data are means ± s.e.m.

**Extended Data Fig. 11.**
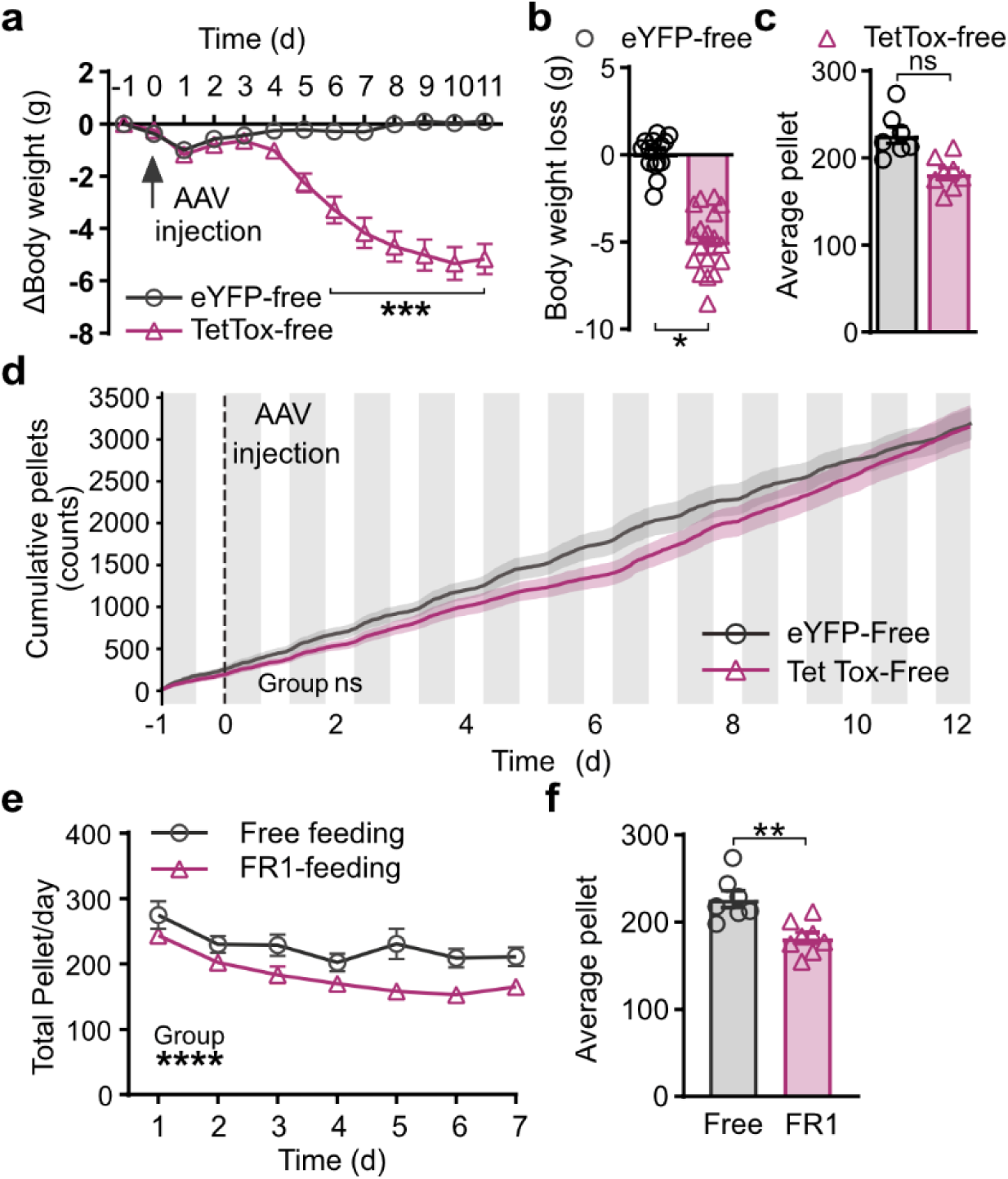
Feeding monitored with FED3 free feed code after silencing POA^KOR+^ neurons. **a**, Body weight changes in mice monitored with free feed code after TetTox injection. **b**, Average body weight changes between control and TetTox treated animals in mice monitored with free feed code. **c**, Average food pellets taken per day monitored with free feed code between control and TetTox treated animals. **d**, Cumulative food intake monitored with FED3 device free feed code between control and TetTox treated mice. **e**, Comparison of pellet taken per day between free feed and FR1 feeding code in FED3 device over a 7-day recording time period. **f**, Comparison of average pellet taken per day between free feed and FR1 feeding code in FED3 device. Two-way ANOVA comparing control and treated groups in (**a**), (**d**), and (**e**). Unpaired-sample *t* test in (**b**), (**c**) and (**f**). *P < 0.05, **P < 0.01, ***P < 0.001, and ****P < 0.0001; ns, no significance. n=7-8 mice per group. Data are means ± s.e.m.

**Extended Data Fig. 12.**
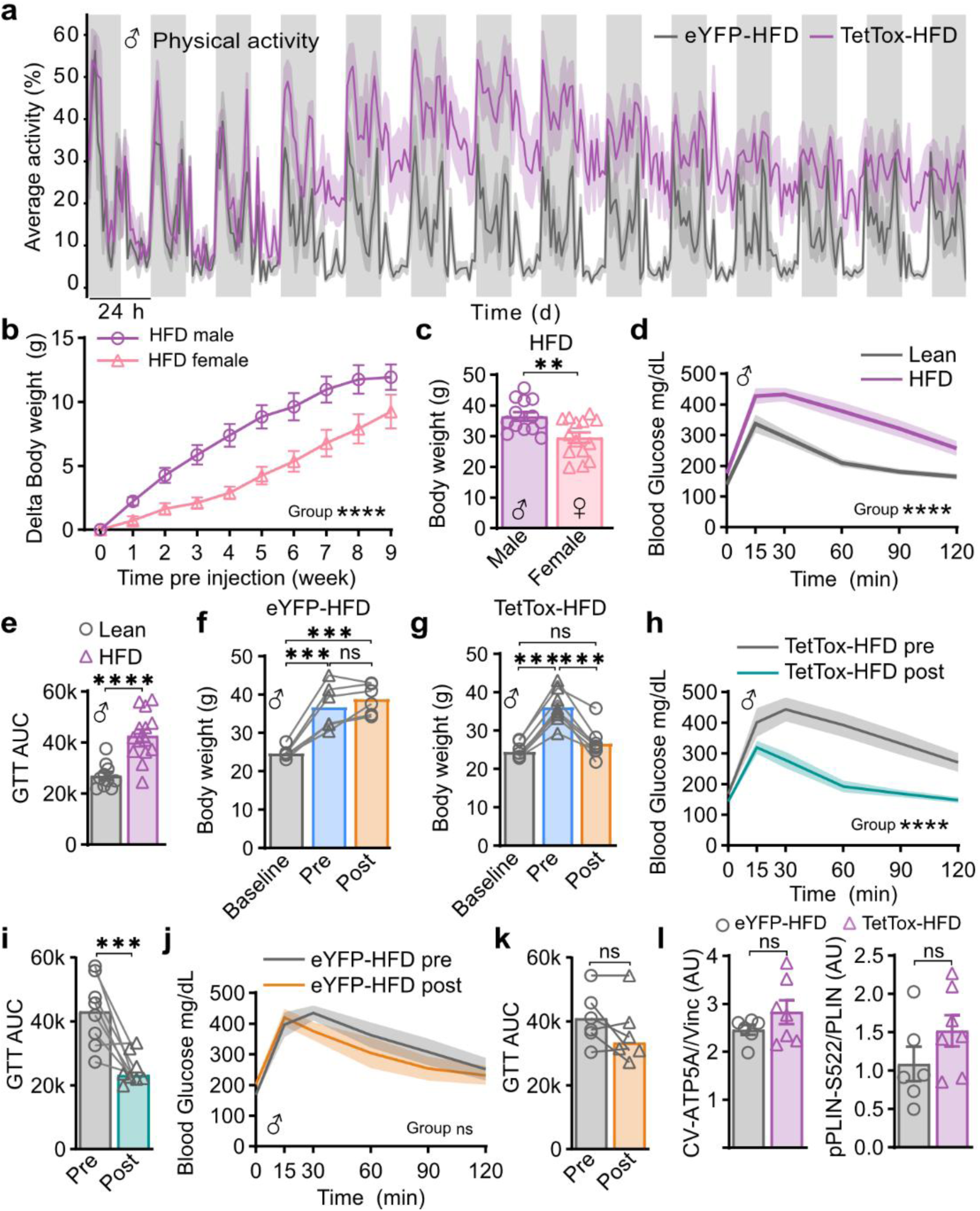
Synaptic inactivation of hypothalamic KOR neurons improves glucose metabolism in diet-induced obese male mice. **a,** Physical activity changes in control and TetTox treated obese mice over two weeks after viral injection. n=6 vs. 8. **b,** Comparison of body weight gain between male and female mice over the 10 weeks HFD feeding before viral injection. n=13 vs. 14. **c,** Comparison of absolute body weight between lean and HFD fed mice right before viral injection at 10 weeks post HFD. n=10-14 per group. **d,** Comparison baseline blood glucose levels during the GTT between lean and HFD fed male mice. n=10-14 per group. **e,** Quantification of the AUC of GTT. **f,** Comparison of body weight changes at baseline, pre and post viral injection of control obese mice. n=6. **g,** Comparison of body weight changes at baseline, pre and post viral injection of TetTox treated obese mice. n=8. **h,** Blood glucose levels during the GTT between pre and post viral injection in TetTox treated obese male mice. n=8. **i,** Quantification of the AUC of GTT. **j,** Blood glucose levels during the GTT between pre and post viral injection in control obese male mice. n=6. **k,** Quantification of the AUC of GTT. **l,** The quantified ratio of protein OXPHOS and PLIN for both groups. Statistical analyses included Two-way ANOVA comparing control and treated groups in (**a**), (**b**), (**d**), (**h**), and (**j**). One-way ANOVA test in (**f**) and (**g**). Unpaired-sample t test in (**c**), (**e**), (**i**), (**k**), and (**l**). *P < 0.05, **P < 0.01, ***P < 0.001, and ****P < 0.0001; ns, no significance. Data are means ± s.e.m.

**Extended Data Fig. 13.**
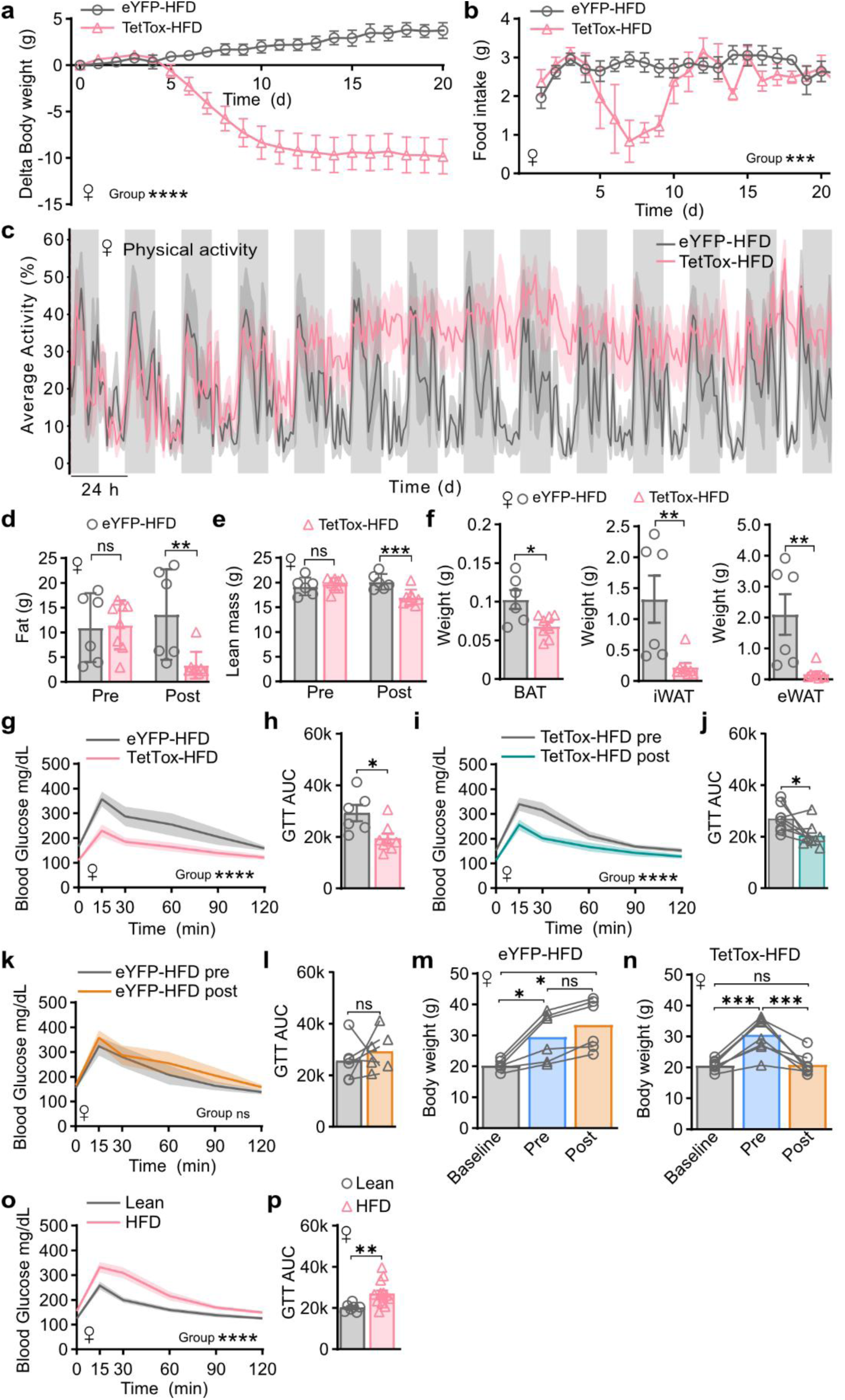
Synaptic inactivation of hypothalamic KOR neurons protects from obesity and ameliorates metabolic syndrome in diet-induced obese female mice. **a**, Body weight changes after silencing POA^KOR+^ neurons showing mice start to lose body weight 5 days after TetTox injection and enter a steady state after two weeks in female mice. **b**, Average food intake per day between control and TetTox treated obese female mice. **c**, Physical activity changes in control and TetTox treated obese female mice over two weeks after viral injection. n=6 vs. 8. **d**, **e**, Fat (**d**) and lean (**e**) masses of control eYFP or TetTox treated obese female mice were measured by EchoMRI at two weeks after viral injection. **f**, Quantification of tissue organ weight of BAT, iWAT, and eWAT after silencing KOR neurons at three weeks after viral injection. **g**, Time course of blood glucose levels during the GTT in control and TetTox treated female mice. **h**, Quantification of the AUC of GTT. **i**, Blood glucose levels during the GTT between pre and post viral injection in TetTox treated obese female mice. n=8. **j**, Quantification of the AUC of GTT. **k**, Blood glucose levels during the GTT between pre and post viral injection in control obese female mice. n=6. **l**, Quantification of the AUC of GTT. **m**, **n**, Comparison of body weight changes at baseline, pre and post viral injection of control (**m**) and TetTox (**n**) obese female mice. n=6-8. **o**, Comparison baseline blood glucose levels during the GTT between lean and HFD fed female mice. n=7-14 per group. **p**, Quantification of the AUC of GTT. Statistical analyses included Two-way ANOVA comparing control and treated groups in (**a**), (**b**), (**c**), (**g**), (**i**), (**k**), and (**o**). One-way ANOVA test in (**m**) and (**n**). Unpaired-sample *t* test in (**d**), (**e**), (**f**), (**h**), (**j**), (**l**), and (**p**). *P < 0.05, **P < 0.01, ***P < 0.001, and ****P < 0.0001; ns, no significance. Data are means ± s.e.m.

## Acknowledgments

The authors acknowledge support through a pilot grant to AJN from The Washington University Nutrition Obesity Research Center (NORC) and the grant P30 DK056341 to the NORC. Studies were also supported by NIH grant R01 NS133365 to ECL. Financial support was also provided by the Department of Anesthesiology at Washington University.

## Author contributions

Conceptualization: AJN, RCL, BNF, JI, ALC

Methodology: AJN, JL, ALC, DF

Formal analysis: ALC, JL, AJN, AE

Investigation: JL, ALC, AJN, AE, HES, DF

Visualization: JL, ALC, AJN, AE, HES

Animal Surgery: JL, ALC, HES, AB

Funding acquisition: ECL, AJN, BNF

Data collection: ALC, JL, HES, BF, AB

Statistical Analysis: AK, JL

Supervision: AJN, BNF, ECL

Writing –original draft: JL, ALC, AJN

Writing – review & editing: JL, ALC, AJN, ECL, BNF

## Additional information

### Declaration of generative AI and AI-assisted technologies in the writing process

During the preparation of this work the author(s) used CHAT-GPT to aid in copy editing proofing the text. After using this tool/service, the authors reviewed and edited the content as needed and take full responsibility for the content of the published article.

